# Characterization of novel competitive inhibitors of *P. falciparum* cGMP-dependent protein kinase

**DOI:** 10.1101/2021.06.24.449800

**Authors:** Tyler Eck, Mariana Laureano de Souza, Rammohan R. Yadav Bheemanaboina, Ramappa Chakrasali, Tamara Kreiss, John J. Siekierka, David P. Rotella, Purnima Bhanot, Nina M. Goodey

**Affiliations:** Department of Chemistry and Biochemistry, Montclair State University, Montclair, NJ 07043, USA; Department of Microbiology, Biochemistry and Molecular Genetics, Rutgers New Jersey Medical School, Newark, NJ 07103

**Keywords:** *P. falciparum*, cGMP-dependent protein kinase, malaria, isoxazole, competitive inhibitor

## Abstract

*P. falciparum* cGMP-dependent protein kinase (PfPKG) is an enticing anti-malarial drug target. Structurally novel isoxazole-based compounds were shown to be ATP competitive inhibitors of PfPKG. Isoxazoles **3** and **5** had *K*_i_ values of 0.7 ± 0.2 and 2.3 ± 0.9 nM, respectively, that are comparable to a known standard, 4-[2-(4-fluorophenyl)-5-(1-methylpiperidine-4-yl)-1H pyrrol-3-yl] pyridine (1.4 ± 0.5 nM). They also exhibited excellent selectivity for PfPKG over the human ortholog and the gatekeeper mutant T618Q PfPKG, which mimics the less accessible binding site of the human ortholog. The human ortholog’s larger binding site volume was predicted to explain the selectivity of the inhibitors for the *P. falciparum* enzyme. Analogs **4** and **6** were at least 20-fold less potent compared to **3** and **5**, suggesting that removing the carbonyl group in **3** or altering the diethylamino moiety in **5** reduced affinity.

## Introduction

Malaria, a mosquito-borne disease caused by the *Plasmodium* parasite, accounted for an estimated 229 million cases and 409,000 deaths in 2019 alone. ^*1*^ Emerging parasite resistance to artemisinin combination therapy threatens the effectiveness of current programs to control the disease and requires the development of new anti-malarials to continue eradication efforts.^*2*^ There is an unmet need for molecules with mechanisms of action that are different from currently-used drugs, as well as drugs that act on multiple parasite stages to provide effective treatment, chemoprotection, chemoprevention and eventual eradication of malaria. ^*2*^ An essential part of the mechanism of action is an understanding of molecular-level interactions between the protein target and ligand. This information can be experimentally derived and/or theoretically produced and often a combination of approaches is useful.

cGMP-dependent protein kinase (PKG) was shown to be a potential chemotherapeutic target in *Plasmodium* and related parasites such as *E. tenella* and *T. gondii*.^*3-6*^ *P. falciparum* PKG (PfPKG) and *P. berghei* PKG function is essential in multiple parasite stages where they control diverse processes, including gametogenesis,^*7*^ merozoite egress and invasion,^*8, 9*^ and late-stage liver development.^*10*^ In accordance with these functions, inhibition of PfPKG blocks parasite infectivity and development.^*3, 7-9*^ PfPKG has a substantially different hydrophobic pocket compared to human PKG (hPKG), a differentiating feature that can provide selectivity.^*4, 5*^ The ‘gatekeeper’ position of PfPKG is occupied by a Thr (Thr618) which has a shorter side chain compared to the Gln at the equivalent position in hPKG. Moreover, human kinases in general have a larger residue, often a Met, at the gatekeeper position.^*11*^ Thus the ‘gatekeeper pocket’ next to the ATP binding pocket is accessible in PfPKG but not in human kinases, making it possible to design inhibitors where a part of the structure occupies the gatekeeper pocket that cannot be accessed in human kinases. These qualities have prompted development of PfPKG inhibitors that can safely treat and/or prevent infection. Baker and co-workers demonstrated success with imidazopyridine **1**, a potent, selective and orally bioavailable PfPKG inhibitor that cleared infection in a mouse model.^*12*^ Its co-crystal structure with *P. vivax* PKG (PvPKG) revealed binding in the ATP pocket and indicated a competitive mode of inhibition of *Plasmodium* PKG. ^*12*^ These observations contributed to an increased emphasis on PfPKG as a potential anti-parasitic drug target.^*12*^ Unfortunately, the imidazopyridine series suffers from genotoxicity issues^*13*^ that substantially limit the value of the chemotype. Characterizing novel chemotypes against PfPKG is one way to address this liability. Characterization against PfPKG of such chemotypes requires a combination of experimental (i.e. mode of inhibition; *in vitro* enzymatic and cell-based assays, assessment of drug-like properties) and theoretical modeling because there are no small molecule-PfPKG crystal structures available.

Our group recently reported the optimization of an isoxazole-based scaffold that lacks any obvious structural safety warnings and demonstrated *in vitro* potency comparable to 4-[2-(4-fluorophenyl)-5-(1-methylpiperidine-4-yl)-1H pyrrol-3-yl] pyridine (**2**) (Figure 1).^*14*^ The trisubstituted pyrrole, **2** was among the first potent inhibitors of PfPKG, blocked development of *P. falciparum in vitro* and *P. berghei in vivo*, and was characterized as an ATP-competitive inhibitor of *E. tenella* PKG.^*4, 5, 8, 9*^ Its use *in vivo* is limited by rapid metabolism to a less active derivative. We believe it is vital to characterize the novel and potentially important series of isoxazole compounds because of the distinct nature of the chemotype, compared to either the imidazopyridine or pyrrole templates. Here we demonstrate the mode of interaction of PfPKG with isoxazole-containing compounds and the structure of two unpublished examples in this series that employ functionality not previously studied in this series. Our data provide insight into the inhibition of PfPKG by this novel chemotype and suggest directions for inhibitor optimization. They also increase the understanding of selectivity determinants in the PfPKG active site and assist the future design of inhibitors that are selective for PfPKG over the human homolog.

**Figure 1.**
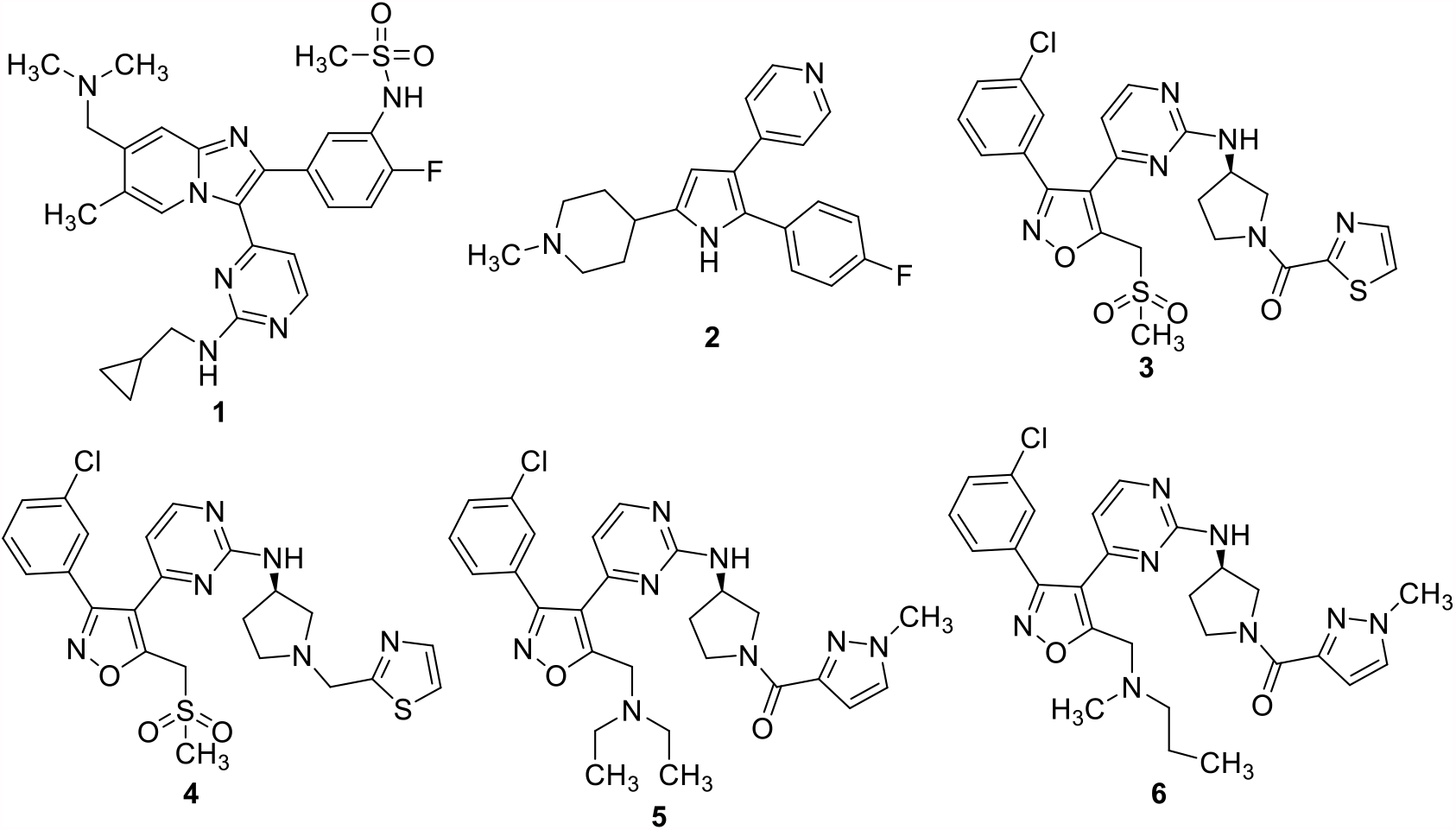
Structures of PfPKG inhibitors. Activities of **2−6** were determined in this work.

## Materials and methods

### Site-directed mutagenesis to generate gatekeeper mutant (T618Q) PfPKG

Site-directed mutagenesis was performed to introduce the T618Q gatekeeper mutation into the full-length PfPKG gene using the following primers: Forward: 5’-CTATTTTCTACAGGAATTAGTAACAGGTGGAG -3; Reverse: 5’-GTTACTAATTCCTGTAGAAAATAGAAATATTTAG -3’. The wild type (WT) PfPKG in the pTrcHis-C vector was a kind gift from the laboratories of David Baker. Reaction conditions included 1X HF buffer (Thermo Fisher Scientific # F530S), ∼100 ng of template DNA, 0.5 μM of each primer, 0.2 mM dNTPs, and 1 unit of Phusion polymerase (Thermo Fisher Scientific # F530S). A PCR protocol was implemented as follows: initial denaturation at 98 °C for 5 minutes; 30 cycles of extension at 98 °C for 30 seconds, 55 °C for 60 seconds, and 72 °C for 3.5 minutes; and a final elongation at 72 °C for 10 minutes. Template DNA was digested with 10 units of DpnI (Fisher Scientific # FERER1701) at 37 °C for 2 hours and the PCR product was transformed into XL Blue cells. The mutation was confirmed by Sanger sequencing.

### Expression and purification of WT and T618Q PfPKG

Full-length WT PfPKG (Uniprot ID Q8I719), a kind gift from David Baker,^*6*^ and T618Q PfPKG were expressed in BL21 (DE3) Star cells. The genes were cloned into the pTrcHisC vector, which incorporates a 6X His-tag at the N-terminus. A 250 mL culture of LB containing 100 µg/mL carbenicillin was inoculated and grown overnight at 37 °C with shaking at 225 RPM. Then 250 mL of fresh media with 100 µg/mL carbenicillin was added and the resulting culture was divided equally into two flasks. IPTG was added to 1 mM and the flasks were incubated with shaking at 18 °C overnight. Cultures were pelleted by centrifugation at 10,000 x g for 15 minutes. The pellets were lysed with 10 mL of ice-cold B-PER extraction reagent (Thermo Fisher Scientific # 78243) containing 1X protease inhibitors (Thermo Fisher Scientific # A32953). Following a 10-minute incubation, the lysis solution was centrifuged at 15,000 x g for 15 minutes and the supernatant was collected.

The soluble lysate was poured over HisPur Cobalt Resin (Thermo Fisher Scientific # 89965) that had been pre-equilibrated with 25 mM HEPES, 20 mM NaCl, 10 mM imidazole pH 7.5. The lysate and resin were incubated, with constant rotation, at 4 °C for 20 minutes. The resin was washed with equilibration buffer until no more protein eluted from the column. WT and T618Q PfPKG were eluted from the column using 25 mM HEPES, 20 mM NaCl, 120 mM KCl, and 250 mM imidazole at pH 7.5. Eluted fractions were tested for the presence of PfPKG by SDS-PAGE and fractions where a protein of 97.5 kDa was detected were collected. Pooled elution fractions were dialyzed in 25 mM HEPES, 20 mM NaCl, 120 mM KCl, and 5% glycerol at pH 7.5. Purified protein was concentrated using a 15-mL Amicon 10 kDa MWCO concentrator (Sigma Aldrich # UFC901024). Enzyme purity and identity were evaluated by SDS-PAGE and Western Blot (Anti-His).

### Expression and purification of hPKG

hPKG (Uniprot ID Q13976) was synthesized by Invitrogen and cloned into the pEF-Bos vector (a kind gift from Dr. Ueli Gubler). The resulting vector was transiently transfected into HEK293 cells using the ExpiFectamine 293 Transfection Kit (Thermo Fisher Scientific # A14525). After enhancement, the culture was expressed for three days at 37 °C with rotation (∼125 RPM) under a 7% CO_2_ atmosphere. The cells containing expressed hPKG were harvested by centrifugation at 10,000 x g for 15 minutes. The resulting pellet was lysed with 10 mL/g of ice-cold M-PER reagent (Thermo Fisher Scientific # 78505) containing 1X protease inhibitor (Thermo Fisher Scientific # A32953). After a 10-minute incubation on ice, the cell lysate was centrifuged at 15,000 x g for 15 minutes. Supernatant was poured over Pierce Glutathione agarose resin (Thermo Fisher Scientific # 16101) equilibrated with equilibration buffer 2 (50 mM Tris, 150 mM NaCl at pH 8) and incubated at 4 °C for 20 minutes. The flow-through was collected and the column was washed with equilibration buffer 2. hPKG was eluted with 50 mM Tris, 150 mM NaCl pH 8 containing 3 mg/mL free glutathione. Enzyme purity and identity were evaluated by SDS-PAGE and Western Blot (Anti-GST).

### Synthesis and characterization of inhibitors

PfPKG inhibitor **2** was synthesized following established protocols.^*15*^ PfPKG inhibitor **5** and **6** were synthesized as previously reported.^*14*^ All reagents were used as provided by the supplier. All reactions were carried out under a nitrogen atmosphere. TLC was carried out on analytical silica gel G254 plates and visualized by UV light. Silica gel chromatography for purification was carried out on a Teledyne Isco Rf200+ using prepacked silica gel columns. NMRs were obtained on a Bruker Avance II instrument. Mass spectra were obtained on an Advion CMS spectrometer and on a Shimadzu LCMS2020 LCMS system. All final compounds were at least 95% pure by NMR and/or LC prior to evaluation. The spectra for these compounds are provided in Supporting Information.

**Scheme 1:**
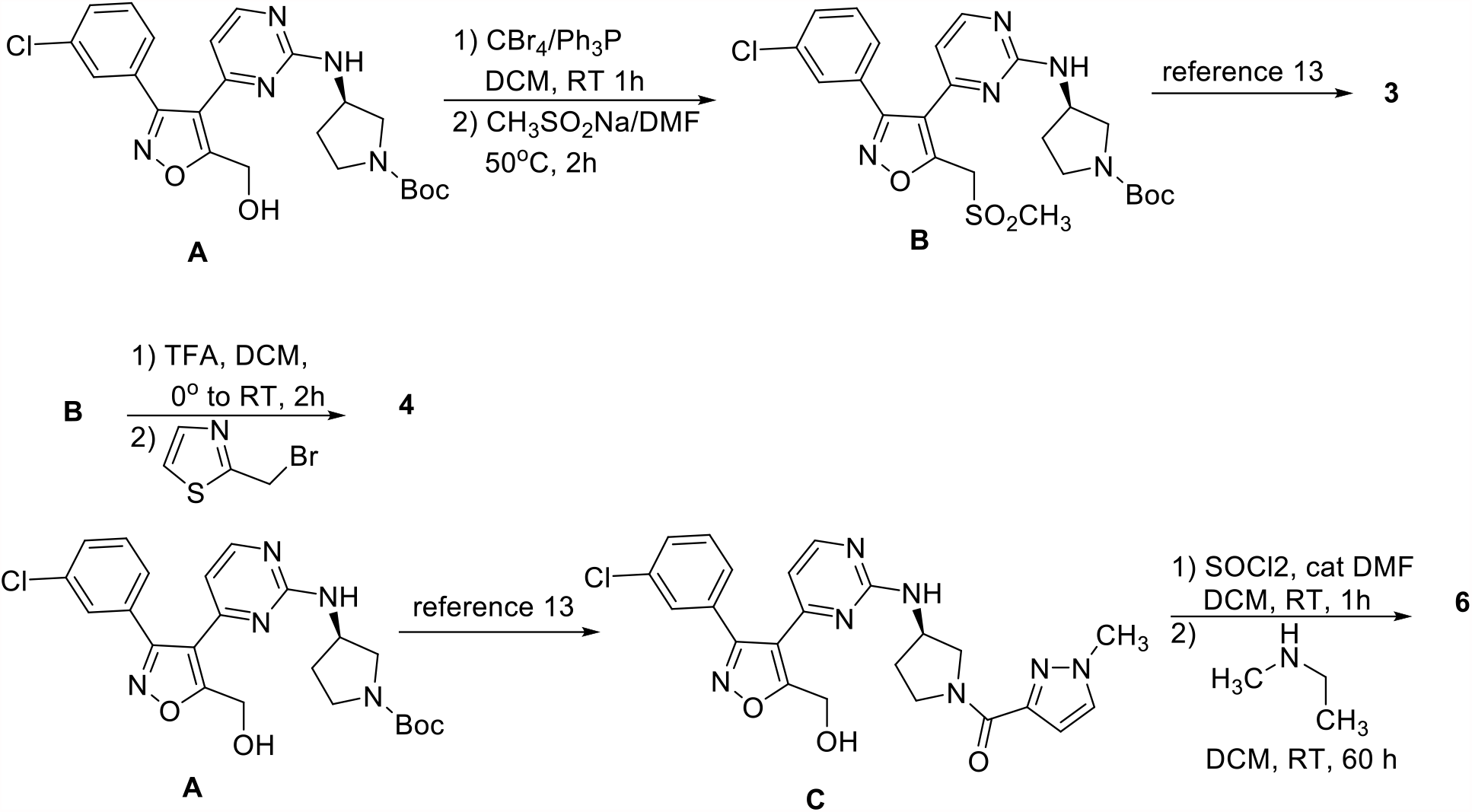
Intermediate **A** and **5** were synthesized as previously reported. ^*14*^

**3**: 1.1 g (2.3 mmol) A was dissolved in 15 mL DCM at room temperature. 1.6 g (4.6 mmol) CBr4 was added, followed by 1.2 g (4.6 mmol) triphenylphosphine. The reaction was stirred at room temperature under a nitrogen atmosphere for 1 hour when TLC indicated the reaction was complete. The reaction was diluted with 100 ml ethyl acetate and washed with 2 × 20 mL water, followed by 2 × 20 mL brine, dried and concentrated. The crude material was purified on a 40g Isco column eluting with 5% methanol in DCM to provide 0.75g (1.4 mmol, 61%) of intermediate **B** as a solid. Boc cleavage and acylation were carried out as previously described. ^*12*^

1H NMR (d, CDCl3, 400 MHz): 8.21 (d, 1H), 7.87 dd, 1H), 7.51 (m, 3H), 7.37 (m, 2H), 6.36 (br s, 1H), 5.76 (br s, 1H), 5.0 (m, 2H), 4.6 (m, 1H), 4.4 (m, 1H), 4.3 (t, 1H), 4.14 (dd, 1H), 3.9 (dd, 1H), 3.83 (m, 1H), 3.09 (d, 3H), 2.33 (m, 1H), 2.1 (m, 1H).

13C (d, CDCl3, 100 MHz): 165, 161, 160, 159.20, 159.5, 143, 134, 130, 129.5, 129.0, 127, 124, 110, 52, 51, 49, 47, 45, 41, 32, 29.

**4**: Intermediate **B** was treated with trifluoroacetic acid in DCM as previously described. ^*14*^ 20 mg (0.046 mmol) of the amine was dissolved in 5 mL DCM and triethylamine (0.138 mmol, 3 equiv.) was added at room temperature. 2-bromomethyl thiazole (10 mg, 0.0552 mmol, 1.2 equiv) was added at room temperature and the reaction stirred for six hours when TLC showed consumption of **B**. The reaction was diluted with 15 mL DCM, washed twice with 20 mL water and brine, dried, concentrated and purified using a 4g Isco column eluting with 5% MeOH in DCM to provide 18 mg (0.034 mmol, 74%) of **4** as a solid.

1H NMR (d, CDCl3, 400 MHz): 8.20 (d, 1H), 7.70 (d, 1H), 7.54 (s, 1H), 7.47 (m, 1H), 7.37 (m, 2H), 7.27 (d, 1H), 6.32 (br s, 1H), 5.61 (d, 1H), 5.01 (apparent q, 2H), 4.55 (m, 1H). 4.03 (d, 2H), 3.08 (m, 4H), 2.80 (m, 2H), 2.54 (apparent q, 1H), 2.40 (m, 1H).

13C (d, CDCl3, 100 MHz): 170, 161, 160, 142, 134, 130.4, 130.1, 129.5, 129.0, 127, 119.6, 119.4, 60, 56, 52.5, 52.1, 50.69, 50.62, 41.

Intermediate **C**: 2.00g (4.23 mmol) intermediate **B** was reacted with TFA in DCM, followed by amide coupling as previously described ^*14*^using 1-methyl pyrazole-3-carboxylic acid to afford 1.6 g (3.34 mmol, 79% yield) as a solid.

1H NMR (d, CDCl3, 400 MHz): 8.10 (m, 1H), 8.0 (m, 1H), 7.80 (m, 1H), 7.0 (m, 4H), 6.52 (br s, 1H), 6.46 (apparent t, 1H), 3.92 (s, 2H), 3.86 (m, 3H), 3.74 (m, 1H), 3.65 (m, 2H), 2.3 (m, 1H), 2.1 (m, 1H)

**6**: 100 mg (0.21 mmol) of intermediate **C** was dissolved in 10 mL DCM and cooled to 0oC in an ice bath. 1 drop of DMF was added, followed by 38 mg (0.32 mmol, 1.5 equiv) thionyl chloride. After 15 minutes, TLC analysis (8% MeOH/DCM) indicated the reaction was complete. The reaction was diluted with 10 mL DCM and washed with water until neutral. The solution was dried and concentrated. The crude chloride was used without further purification. 50 mg (0.10 mmol) of this material was dissolved in 5 mL DCM at room temperature. 37 mg (0.50 mmol) of N-methyl propyl amine was added and the reaction was stirred at room temperature for 60 hours when TLC (10% MeOH/DCM) indicated consumption of starting material. The reaction was diluted with 20 mL DCM and washed twice with 10 mL brine. The crude product was purified using a 4 g Isco prepacked column eluting with 10% MeOH/DCM to afford 30 mg (0.056 mmol, 56%) of the desired product as a solid.

1H NMR (d, CD3OD, 400 MHz): 8.35 (apparent t, 1H), 8.08 (d, 1H), 7.84 (d, 1H), 7.44 (m, 4H), 6.5 (br s, 1H), 4.4 (m 3H), 3.5-3.9 (m, 7H), 2.45-2.72 (m, 5H), 2.3 (m, 1H), 2.1 (m, 1H), 1.59 (m, 2H), 0.9 (m, 3H)

### IMAP assay to determine PfPKG and hPKG specific activities

WT and mutant PfPKG kinase activities were determined using the commercial immobilized metal ion affinity-based fluorescence polarization (IMAP) assay (Molecular Devices # R8127).^*16*^ The kinase assay wells (20 μL total volume) contained assay buffer RB-T (10 mM Tris-HCl, pH 7.2, 10 mM MgCl_2_, 0.05% NaN_3_, 0.01% Tween®20), recombinant PfPKG (ranging from ∼0.2 – 0.002 mg/mL in the well), 120 nM fluorescent peptide substrate (FAM-PKAtide), 10 μM ATP, 1 μM cGMP, and 1.0 mM DTT. The reactions were initiated by the addition of FAM-PKAtide and incubated for one hour. Then 60 μL of the Progressive Binding Reagent (PBR) mixture was added and the resulting solutions (80 μL total volume) were incubated for 30 minutes. The PBR mixture was made according to the commercial protocol for FAM-PKAtide substrate (100% 1X IMAP Progressive Binding Buffer A combined with PBR diluted 400-fold). Fluorescent polarization was read parallel and perpendicular to the excitation plane (ex. 485 nm/ em. 528 nm) using a Synergy 2 Microplate reader (BioTek, Winooski, VT) and the relationship between signal and time was linear (Figure S2). The averages of the signals from each experimental well were calculated (n=2). Various PfPKG concentrations were tested and the resulting signals (mPolarization) were graphed against total enzyme concentrations used in the well (mg/mL). The slopes of these graphs corresponded to the PfPKG specific activities and were used to find initial screening conditions and to ensure consistency of purification quality. The specific activity of hPKG was tested using a similar protocol with the following modifications: The substrate FAM-IP3R-derived peptide (RP7035) and the corresponding commercial protocol for the PBR mixture (75% 1X IMAP Progressive Binding Buffer A, 25% 1X IMAP Progressive Binding Buffer B, PBR diluted 600-fold) were used. After addition of the PBR mixture, the solutions were incubated for 1 hour instead of 30 minutes.

### Inhibitor IC_50_ determination

IC_50_ values were determined using the IMAP assay conditions described above. Inhibitor concentrations ranged from 4 nM to 10 µM in the wells. The following amounts of enzymes were used in each assay well: 9 nM of WT PfPKG, 4 nM of T618Q PfPKG, and 5 nM of hPKG (to achieve a maximal signal of ∼250 mPolarization units). Additionally, enzyme was preincubated with inhibitor concentrations for 15 minutes at room temperature prior to adding the enzyme-inhibitor solution into the assay wells. The trisubstituted pyrrole (**2**) was used as a positive control in each experiment. The data were analyzed using a four-parameter logistic curve using Microsoft Excel Solver and dose response curves were generated using Microsoft Excel.

### Kinetic IMAP assay for determination of Michaelis constant (K_m_) of ATP and FAM-PKAtide

The IMAP assay was adapted to a kinetic format to determine initial velocities and Michaelis constants for ATP for the PfPKG enzymes. Instead of incubating the reaction for 1 hour as described above, the reaction was allowed to proceed for different time periods ranging from 0 to 70 minutes. The kinetic IMAP assay was implemented by initiating the 20-minute incubation period of the fluorescent peptide, cGMP, inhibitor and PfPKG at different times, starting with the wells that required the longest incubation. After a well had been incubated for 20 minutes, ATP was added to the well to initiate the reaction (this was done at different times depending on the desired reaction time, starting with the wells that required the longest incubation times). The reaction mixtures were then incubated for desired amounts of time and the PBR developing solution was added to all reactions at once, stopping all reactions at the same time. After the final incubation, fluorescence polarization was read and velocities were determined from the change in polarization over time using data points where the relationship between polarization and time was linear (at least 60 minutes) (Figure S2). Velocities were determined at different concentrations of ATP (0.78 – 100 µM) while keeping the FAM-PKAtide and cGMP at original concentrations described above. Enzyme amounts used for these experiments were 28 nM of WT PfPKG and 12 nM of T618Q PfPKG in the well. Initial velocities were converted to percent activities and plotted against substrate concentrations in KaleidaGraph. The resulting curves were fit to the Michaelis-Menten equation. Reported *K*_m_ values were the result of three replicate measurements for WT PfPKG and four measurements for T618Q PfPKG. Statistical significances were evaluated using a two-sided Student’s t-test.

### Inhibitor K_i_ and mechanism of inhibition determination by Dixon plots

The mechanism of inhibition was kinetically determined by measuring reaction rates in the presence of varying concentrations of inhibitor and substrate. The procedure was conducted as described above for determining PfPKG *K*_m_ values, with a few alterations. Namely, inhibitors were pre-incubated with PfPKG, cGMP, and FAM-PKAtide for 20 minutes before the reaction was initiated with ATP (Final ATP Concentrations in the well: 12.5, 25, 37.5, and 50 µM). Four novel isoxazole inhibitors **(3-6**) and the trisubstituted pyrrole **2** were tested by this method. The concentrations of inhibitors in assay wells were as follows: **2** (2.5 and 5 nM), **3** (2.5 and 5 nM), **4** (250 and 500 nM), **5** (2.5 and 5 nM), and **6** (100 and 200 nM). Velocities for each combination of inhibitor and ATP concentrations were determined as described above and the reciprocal velocities were plotted against inhibitor concentration in Excel. The data were fitted to linear equations for each substrate concentration. The average of the inhibitor concentrations at the intersection points of all lines was determined and corresponded to the -*K*_i_ value for that experiment. The reported *K*_i_ values were the result of three replicate measurements.

### Molecular docking simulations

The PfPKG structure (PDB ID 5DYK)^*17*^ and hPKG structure (PDB ID 6BDL) were prepared by removing water molecules and adding hydrogen atoms in GOLD 5.8.1 (Cambridge Crystallographic Data Centre, Cambridge, UK).^*18*^ The PfPKG mutant T618Q structure was generated by homology modeling in the Molecular Operating Environment (MOE) (Chemical Computing Group, Montreal, CA)^*19*^ using the PfPKG structure (PDB ID 5DYK) as a model. MOE was used for the necessary simulations to prepare the mutant protein structure before docking. The optimal model is provided based on Boltzmann-weighted randomized samples of backbone and side chain conformations, scored based on contact energy function.^*20*^ The three dimensional ligand coordinate .sdf files for **3-6** were generated in ChemDraw. Molecules **3-6** were docked with the program GOLD 5.8.1 against PfPKG, PfPKG mutant T618Q, and hPKG. Default parameters of GOLD, that include a 10 Å radius, were used for docking and the generated poses were evaluated using the GoldScore function with the exception that the search efficiency for the genetic algorithm was increased to 200%. The binding pocket was defined as a sphere, with a radius of 10 Å, centered on the coordinates of the Thr618 oxygen atom in the WT PfPKG. In the T618Q mutant PfPKG the binding pocket was centered around the Gln618 oxygen atom and around the Met438 sulfur atom in the hPKG. These results were compared to those using a sphere with a 20 Å radius centered on the same coordinates.

## Results and Discussion

The IC_50_ of **2** and four isoxazole compounds (**3-6**) against recombinant, wild type (WT) PfPKG were determined using FAM-PKAtide as substrate in an IMAP assay^*16*^ (Table 1, Figure S1-S3). The IC_50_ of reference compound **2** was determined to be 31 ± 6 nM (Table 1). This is higher but qualitatively similar to the previously reported IC_50_ (8.53 nM) against partially purified PfPKG obtained using a ^33^P-phosphorylated peptide substrate.^*3*^ The IC_50_ of **3** and **5** were found to be 14 ± 1 and 21 ± 11 nM, respectively, and similar to that of **2**. On the other hand, **4** and **6** were nearly 57-fold and 20-fold less potent, respectively than their matched pair analogs, **3** and **5**. The interactions lost or altered from removing the carbonyl in **3** or altering the diethylamino moiety in **5** significantly reduced binding affinity (p < 0.05) (Figure 1).

**Table 1.** Experimental determination of IC_50_ and *K*_i_ of **2**-**6** for WT PfPKG. Estimated *K*_i_ was calculated using the tight-binding Cheng-Prusoff Equation.^*21-23*^ Experimental Ki was determined by Dixon Plot. Values are the mean of at least two biological replicates ± standard error.

| Compound | $IC_{50}^{WT}$ (PfPKG)<br>(nM) | Estimated $K_i$<br>(nM) | Experimental $K_i$<br>(nM) |
| --- | --- | --- | --- |
| **2** | $31 \pm 6$ | $1.7 \pm 0.4$ | $1.4 \pm 0.5$ |
| **3** | $14 \pm 1$ | $0.70 \pm 0.04$ | $0.7 \pm 0.2$ |
| **4** | $792 \pm 115$ | $50.8 \pm 7.4$ | $210 \pm 41$ |
| **5** | $21 \pm 11$ | $1.1 \pm 0.7$ | $2.3 \pm 0.9$ |
| **6** | $425 \pm 133$ | $27.1 \pm 8.5$ | $87 \pm 24$ |

While **2** was demonstrated to compete with ATP in *E. tenella* PKG,^*5*^ the mechanisms of PfPKG inhibition by **2** or the structurally distinct isoxazoles have not been empirically determined. In order to investigate their binding affinities and mechanism of action, we adapted the IMAP assay to a kinetic format (see Methods for details). The IMAP assay eliminates the need for radioactive substrates and ATP and can be implemented quickly in a 96-well assay format. Rather than reading a single, quantitative reaction endpoint with the traditional IMAP protocol, we measured fluorescence polarization at different time points and used the change in polarization over time to determine reaction velocities. This made it possible to determine reaction kinetics and to calculate *K*_m_ and *K*_i_. To our knowledge, this is the first reported modification of the simple mix and read IMAP assay, developed by Molecular Devices, for use in kinetic measurements.

Using the kinetic IMAP assay, we generated Dixon plots and determined inhibition constants (*K*_i_) for **2**-**6**.^*21*^ Figure 2 shows the inverse rates of WT PfPKG at various concentrations of ATP (12.5, 25, 37.5 and 50 µM), and **2** or **3** (0, 2.5 and 5 nM). The intersection points of the lines were found in the second quadrant and indicated that both compounds are ATP-competitive inhibitors of PfPKG. We found this to be characteristic for all compounds tested (Figure S4). *K*_i_ obtained experimentally for each compound are listed in Table 1.

**Figure 2.**
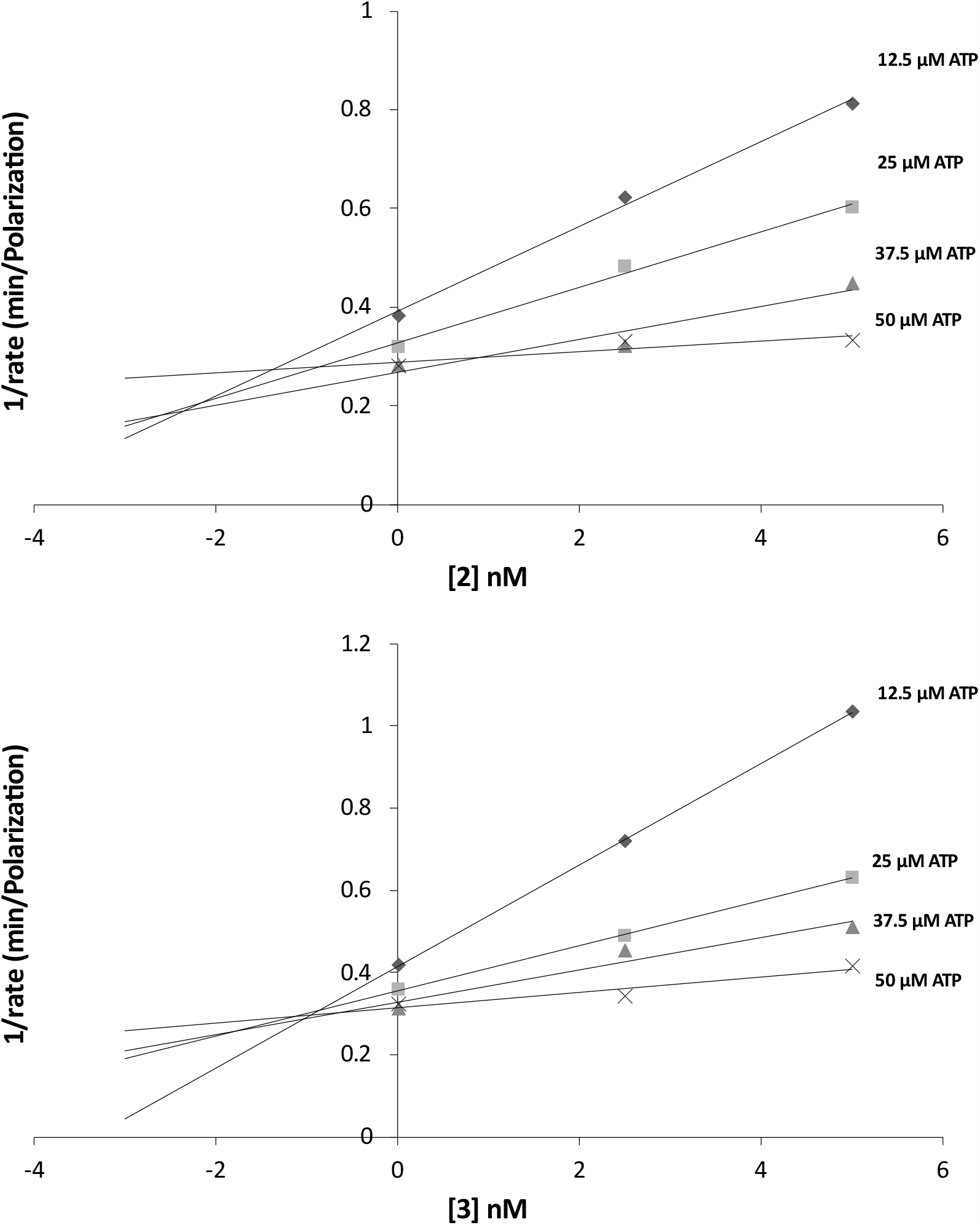
Experimental determination of Ki. A Dixon Plot showed inverse initial velocities in the presence of **2** and **3** at different ATP concentrations. The intersection points for both graphs in the top left quadrant corresponded to a mechanism of competitive inhibition. Data shown are representative of three biological replicates used to determine means reported in Table 1.

We determined the Michaelis constant (*K*_m_) of WT PfPKG for ATP is 6.9 ± 0.1 µM and for T618Q is 10.0 ± 1.6 µM (Figure 3). It is similar to the *K*_m_ of *E. tenella* PKG for ATP (12 ± 2 µM) determined using [γ-^33^P]ATP and the peptide substrate (biotinyl-*e*-aminocaproyl-GRTGRRNSI-OH).^*5*^ The *K*m of PfPKG was used to convert IC_50_ of **3-6** (Table 1) to estimated *K*i, using the Cheng-Prusoff equation.^*21*^ We found our experimentally determined *K*_i_’s to be in agreement with the estimated values that were predicted by the Cheng-Prusoff equation.^*22, 23*^ Our experimental *K*_i_ data showed that **3** (*K*_i_ : 0.7 ± 0.2 nM) and **5** (*K*_i_ : 2.3 ± 0.9 nM) had similar affinities for PfPKG as (Ki: 1.4 ± 0.5 nM). As expected, the two less potent compounds based on IC_50_ values, **4** (*K*_i_ : 210 ± 41 nM) and **6** (*K*_i_ : 87 ± 24 nM), had ∼2 orders of magnitude higher *K*_i_ values compared to **2, 3** and **5**.

**Figure 3.**
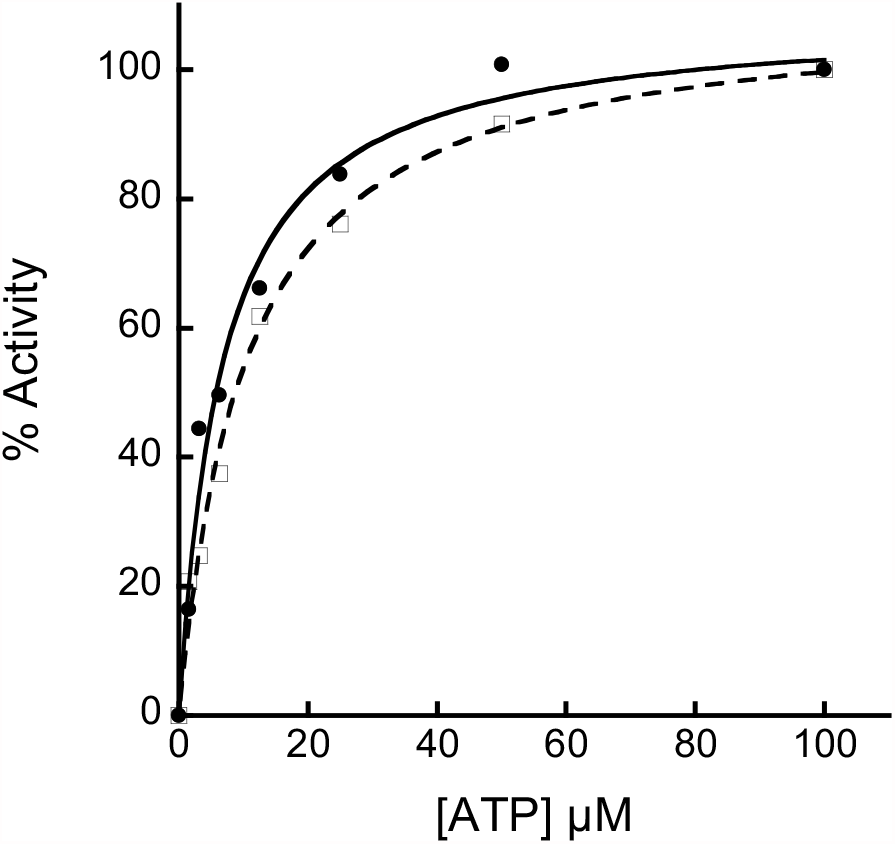
Experimental determination of K_m_. Percent activities of WT (•) and mutant T618Q PfPKG (□) with various ATP concentrations. Data were fit to the Michaelis-Menten equation using KaleidaGraph. Data shown are representative of three biological replicates used to determine the mean.

In order to develop hypotheses to explain the relative binding affinities of structurally-similar pairs of compounds, **3** versus **4**, and **5** versus **6**, we conducted molecular docking studies. We tested binding pockets defined by radii of 10 Å, 20 Å and 30 Å. The same protein-molecule interactions were observed in all cases (data not shown). Therefore, further docking studies were conducted using a binding pocket of 10 Å, the default parameter in GOLD. Evaluation criteria for selection of the preferred pose for each molecule were based on previously observed and predicted ligand interactions with PfPKG and PvPKG and experimentally determined *K*_i_ values in our study. Docking energy scores were not sufficiently different from each other to be used for pose selection.

Previous crystallographic and docking studies have examined binding of three different ATP-competitive inhibitors with *P. vivax* PKG (PvPKG) or PfPKG (for example, PvPKG bound to ML-10 (PDB: 5EZR)).^*12*^ This structure shows ligand interactions with Val614 (Val621 in PfPKG), Thr611 (Thr618 in PfPKG), Asp675 (Asp682 in PfPKG) and Phe676 (Phe683 in PfPKG). ^*12*^ Modeling interactions between PfPKG and a trisubstituted thiazole using the apo PfPKG crystal structure (PDB: 5DYK) identified ligand interactions with Thr618, Asp682, Val621 of PfPKG, as well. ^*24*^ Additional interactions were predicted with Lys570 and Tyr822. Similarly, modeling interactions between PfPKG and a trisubstituted imidazole identified Asp682 and Val621 as critical interactions.^*25*^ In addition, this modeling study suggested an interaction with Glu625. Together, three studies^*12, 24, 25*^identified Asp682 and Val621 as forming critical interactions with ATP-competitive inhibitors of PfPKG. We used these residues as one factor to guide pose selection in our modeling studies based on the assumption that residues described previously, to interact with ligands that occupy the ATP-binding site, are more likely to be involved in interactions with **3, 4, 5**, and **6**. Below we describe the use of the docking results to develop hypotheses about molecular interactions that may be responsible for the different binding affinities of **3, 4, 5**, and **6**.

Since **3** has a lower experimentally determined *K*_i_ than **4**, it is predicted to have stronger and/or more interactions than **4**. The pose selected for the PfPKG-**3** complex (Figures 4C and 4E) located in a similar position as the ATP analog phosphoamino phosphonic acid-adenylate ester (AMPPNP) in an experimentally determined complex with PvPKG (5DZC). ^*17*^ This predicted model is consistent with kinetic data which indicated that **3** is an ATP competitive inhibitor of PfPKG. Compound **3** has a *K*_i_ value approximately 300-fold smaller compared to **4**. The only structural difference between **3** and **4** is a carbonyl group (indicated by an arrow in Figures 4A and B). The selected PfPKG-**3** pose showed five interactions between PfPKG and **3**. Three of these interactions were also present in the selected pose of **4**, including an arene-arene interaction with Phe616, a hydrogen bond with the backbone of Val621, and a hydrogen bond with the sidechain of Thr593 (Figures 4D and 4F). The higher binding affinity of **3** is likely driven by hydrogen bonding between its carbonyl group and the backbone amide of Asp682 in PfPKG. In addition, the pyrimidine group of **3** showed a hydrogen bond with Lys570 in the selected pose. We hypothesize that the lower binding affinity of **4** may be attributed, in part, to the absence of the corresponding carbonyl group and the subsequent lack of this key hydrogen bond with Asp682. Compound **4’s** interaction with Val555 in the selected pose was a van der Waal’s interaction (arene-H), which is expected to be weaker than Lys570’s side chain hydrogen bond with **3**, and this difference could contribute to **4**’s lower binding affinity.

**Figure 4.**
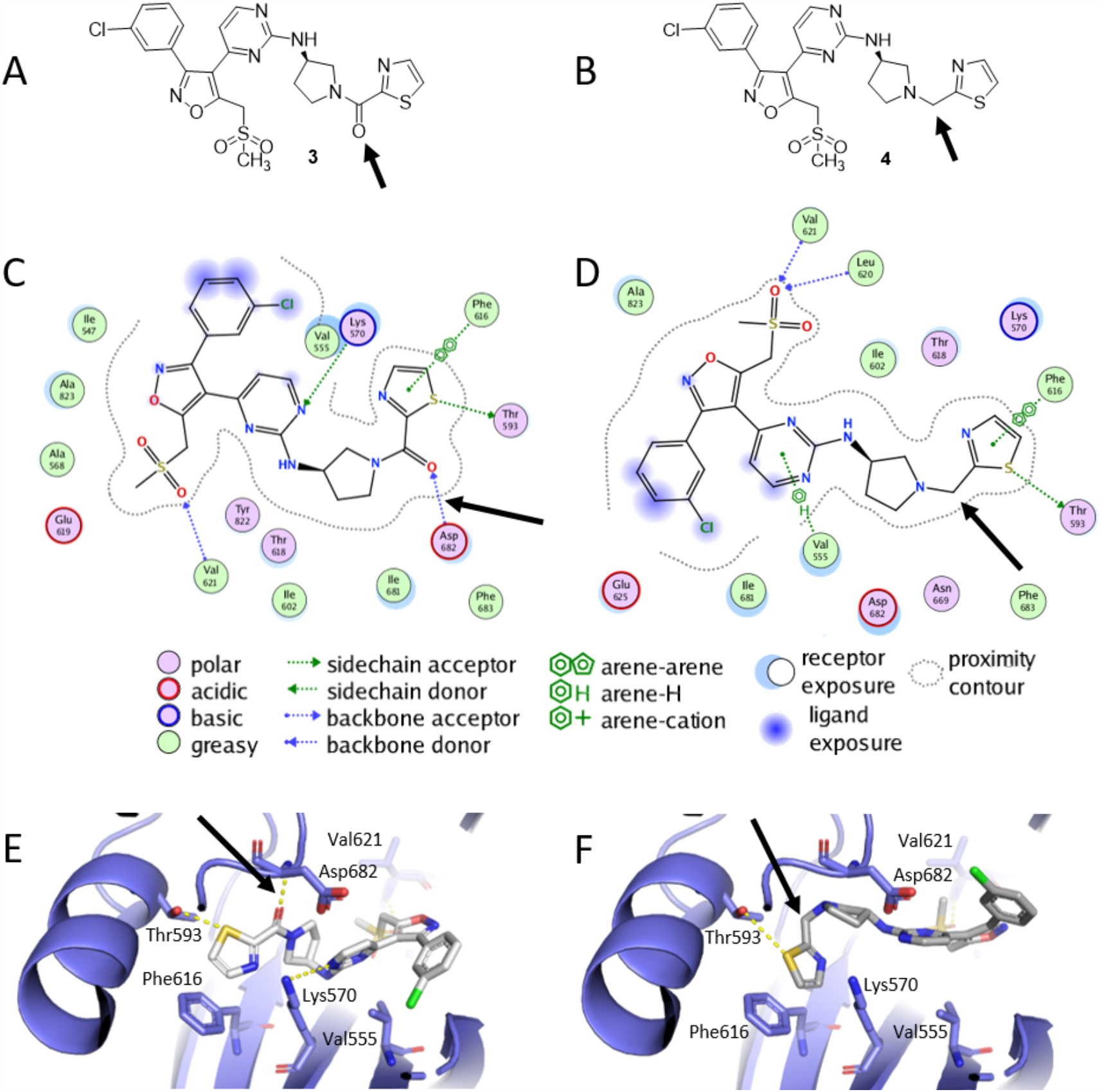
PfPKG interactions with **3** and **4**. Two-dimensional structures of **3** (A) and **4** (B) are shown with their structural difference indicated by a black arrow. PfPKG interactions with **3** (C) and **4** (D) are shown as 2D cartoons with differences indicated by black arrows. Binding of **3** (E) and **4** (F) to PfPKG is depicted in 3D. PfPKG amino acid residues (carbons in blue) and **3** and **4** (carbons in white) are illustrated as sticks. Intermolecular interactions are displayed as dashed yellow lines. Figures C and D were prepared using MOE.^*19*^ Figures E and F were prepared using PyMOL.^*26*^

The *K*_i_ value of **5** was ∼38-fold lower for PfPKG than **6** indicating that **5** has a higher binding affinity. The structural difference between **5** and **6** is in the amino substituent (indicated by an arrow in Figure 5A and B). Both compounds are tertiary amines; **5** is symmetrically substituted while **6** contains methyl and n-propyl groups, and their *K*_i_ data indicate a steric limitation in this region. The selected docking poses showed the two compounds adopting different conformations inside the binding site. In **5**, the diethylamino group and in **6**, the chlorobenzyl group was exposed to solvent. The isoxazole rings of both compounds formed arene-cation interactions with the Lys570 side chain. Docking predictions suggested van der Waal’s (arene-H) interactions between the pyrazole group of **5** and Tyr822 and the pyrazole group of **6** and Ala823. In the selected pose for the PfPKG-**5** complex, the backbone carbonyl of Glu619 formed a hydrogen bond with an amine in **5**. This hydrogen bond was absent in the selected pose for PfPKG-**6** complex (black arrow in Figures 5C and 5D). We hypothesize that hydrogen bonding with Glu619 is the likely driver of **5**’s lower Ki compared to **6**. Alternative poses for 5 are possible but were less favored because they did not provide sufficient explanation for **5**’s higher binding affinity for PfPKG compared to **6**.

**Figure 5.**
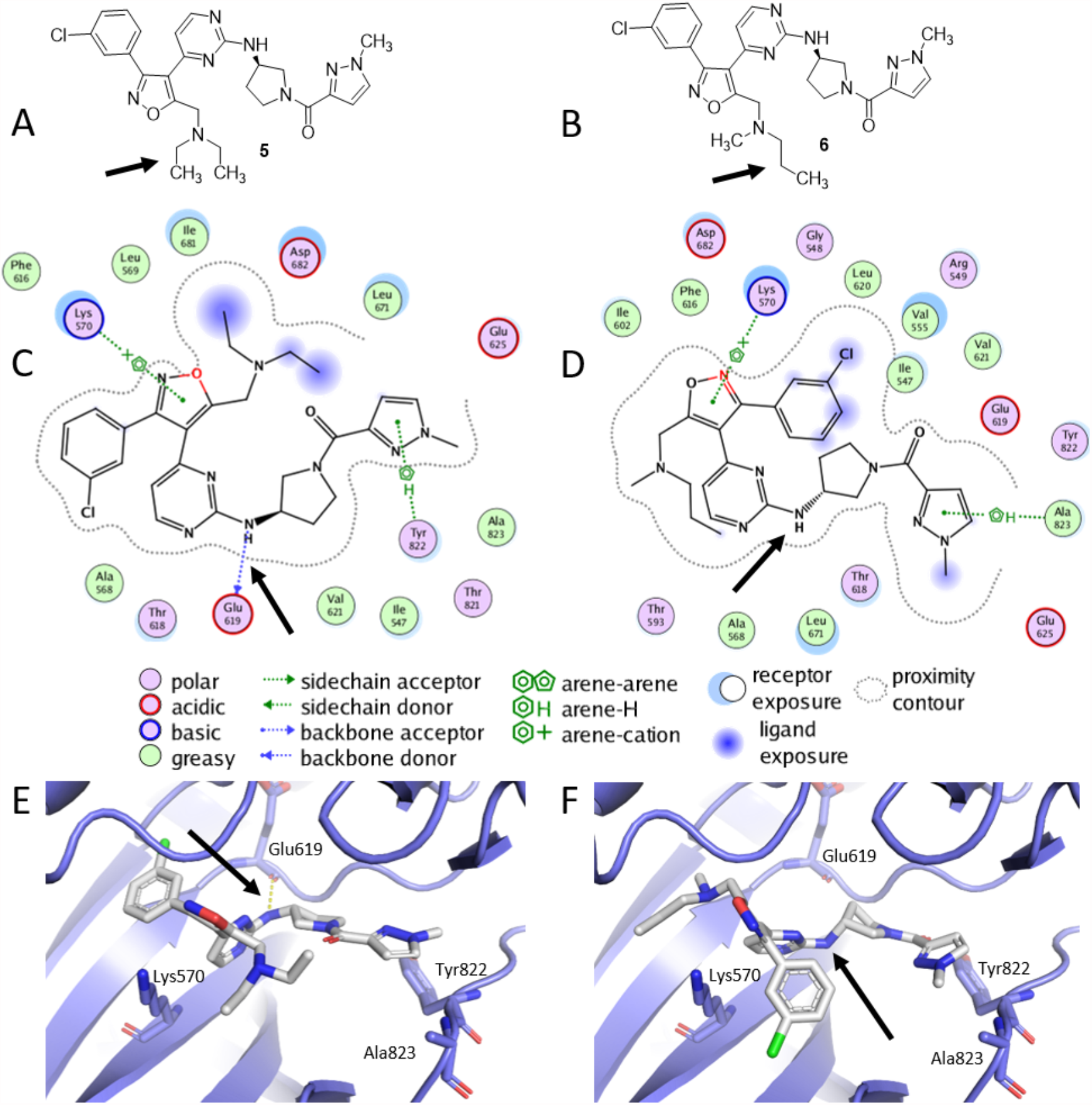
PfPKG interactions with **5** and **6**. Two-dimensional structures of **5** (A) and **6** (B), with their structural differences indicated by black arrows. PfPKG interactions with **5** (C) and **6** (D) are shown as 2D cartoons, with differences indicated by black arrows. Binding mode of **5** (E) and **6** (F) in PfPKG are depicted in 3D. PfPKG amino acid residues (carbons in blue) and **5** and **6** (carbons in white) are illustrated as sticks. Intermolecular interactions are displayed as dashed yellow lines. Figures C and D were prepared using MOE.^*19*^ Figures E and F were prepared using PyMOL.^*26*^

To assess selectivity of compounds for PfPKG over the human enzyme, each molecule was screened for inhibition against hPKG at 10 µM. All compounds were weak inhibitors of hPKG at 10 µM, suggesting excellent selectivity of **2-6** for WT PfPKG over the human enzyme (Table 2). Compounds were also evaluated against the T618Q PfPKG ‘gatekeeper’ mutant, in which access to the hydrophobic pocket appears blocked by the larger side chain of the glutamine substituent (Figures S5 and S6). At 10 µM, **2-6** were inactive or weak inhibitors of T618Q PfPKG. The T618Q PfPKG mutant was found to have a small but significant difference in *K*_m_ for ATP (10.0 ± 1.6 µM) compared to the WT PfPKG (6.9 ± 0.1 µM) (*p* = 0.027) (Figure 3). Previous work similarly found that the *K*_m_ of the *E. tenella* PKG gatekeeper mutant was higher than that of the WT enzyme.^*13, 27*^

**Table 2.** Inhibition of hPKG and T618Q PfPKG. Values are the result of at least two replicate measurements and standard errors are shown.

| Compound | % Inhibition at 10 μM<br>(hPKG) | % Inhibition at 10 μM<br>(T618Q PfPKG) |
| --- | --- | --- |
| **2** | $9.0 \pm 7.5$ | $16.0 \pm 4.5$ |
| **3** | $7.4 \pm 0.6$ | $33 \pm 9$ |
| **4** | $13 \pm 4$ | $0 \pm 0.5$ |
| **5** | $19 \pm 9$ | $36 \pm 5$ |
| **6** | $12.2 \pm 0.8$ | $6.5 \pm 8.0$ |

We used the Protein Pocket Volume application in MOE^*19*^ to calculate the difference in volumes of the binding site in WT and T618Q enzymes. The estimated pocket size of WT was found to be 17.1 Å^3^ (13.8%) larger than that of T618Q (Figure S7). In order to obtain insights into **3**’s relative lack of inhibition of T618Q PfPKG, **3** was docked into the binding pocket of T618Q PfPKG (Figure 6). Since **3** was in close proximity to the Thr618 side chain in the WT enzyme, we hypothesize that the larger Gln sidechain in the T618Q mutant pushes **3** into a different binding mode. In addition, fewer interactions were observed when **3** was docked into T618Q PfPKG compared to WT (Figure 6 and Figure S7). For example, the arene-arene interaction between Phe616 and the thiazole ring of **3 (**in the selected pose) was observed in PfPKG-**3** complex but was missing in the T618Q PfPKG-**3** complex (Fig. 6B-D). Similar results, demonstrating fewer interactions with T618Q PfPKG compared to PfPKG, were obtained for **4** (Figure S8). Finally, in order to generate hypotheses aimed at understanding the selectivity of **3** for PfPKG over hPKG, **3** was docked into hPKGα. The selected pose showed **3** retaining only hydrophobic interactions with hPKGα (Fig. 6E-F) and losing atomic interactions that were observed with PfPKG (Fig. 6). These lost interactions may explain why **3** lacks activity against hPKG.

**Figure 6.**
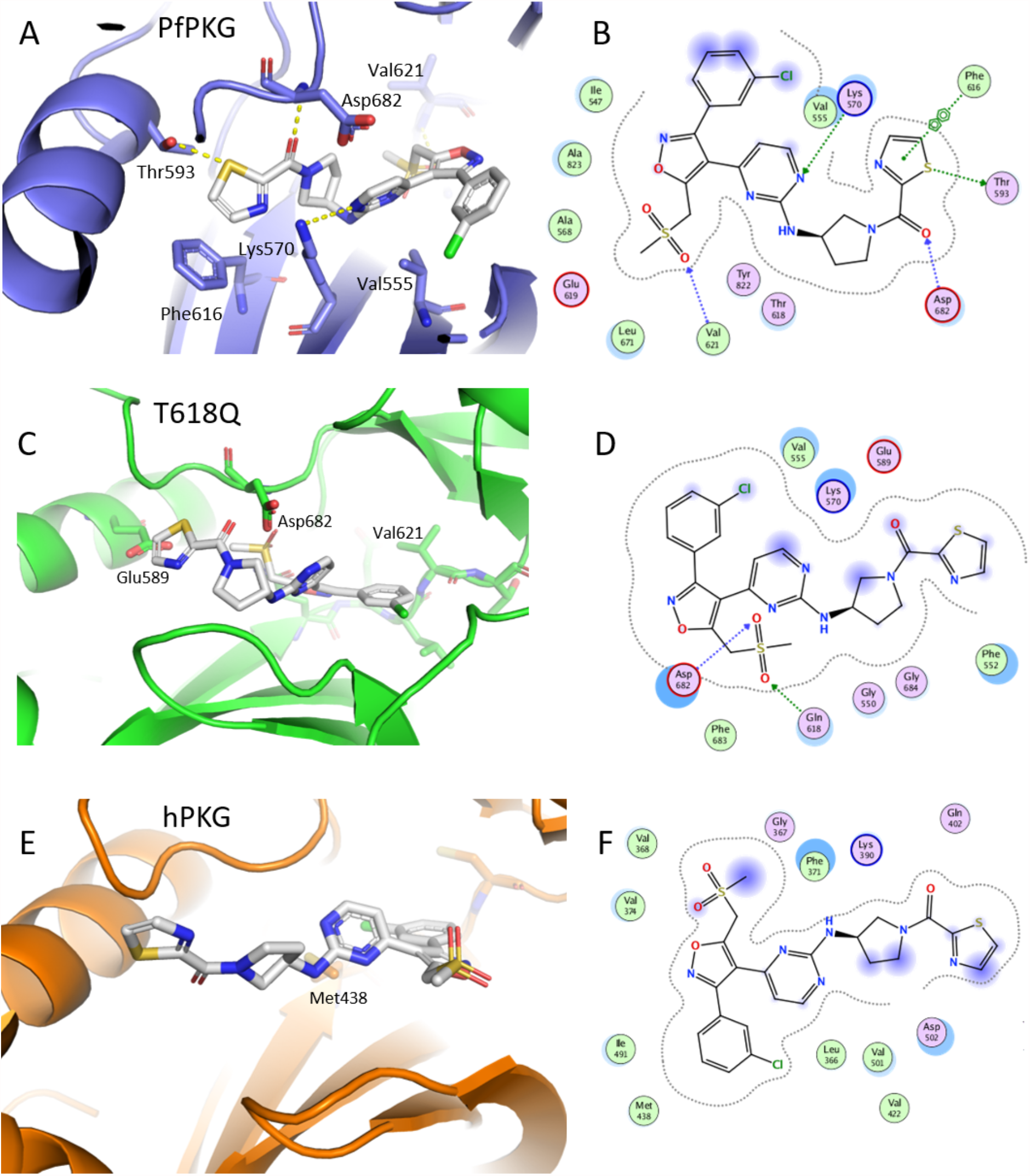
Interactions between **3** and PfPKG, T618Q and hPKG. Interactions in selected poses of **3** docked into WT (A, B), T618Q (C, D) and hPKG (E and F). Figures A, C and E were prepared with PyMOL.^*26*^ Figures B, D and F were prepared with MOE.^*19*^

## Conclusion

In this study, isoxazoles **3−6** were tested as potential inhibitors of PfPKG, T618Q PfPKG, and hPKG. A larger survey of the structure-activity relationships that led to the new compounds reported in this paper was reported previously.^*14*^ While **5** was known to inhibit PfPKG,^*14*^ its mechanism of inhibition was unknown. Compounds **3** and **4** are reported here for the first time. Inhibitory activities of **3−6** against PfPKG and the potential mechanisms of inhibition were investigated, and Dixon plot analyses were used to determine *K*_*i*_ values. Lastly, to generate hypotheses to explain their relative *K*_*i*_ values, **3−6** were docked into the binding pockets of WT PfPKG, the ‘gatekeeper’ PfPKG mutant T618Q, and hPKG. The kinetic data demonstrated for the first time that **2** and structurally novel isoxazoles **3−6** are ATP competitive inhibitors of PfPKG. Compounds **3** and **5** have IC_50_ and *K*_i_ values comparable to **2**. Both **3** and **5** were also highly selective for PfPKG, with low activity against hPKG and T618Q PfPKG. This biochemical information is essential because, when coupled with computational results, it facilitates focused optimization to furnish potent, selective, orally bioavailable and drug-like PfPKG inhibitors that have advantages compared to known chemotypes. Assessment of the drug-like properties and cell-based activity of this series will be the subject of future publications.

Computational docking studies with PfPKG combined with biochemical results obtained with **3−6** made it possible to form reasonable structure-based hypotheses for affinity changes detected by kinetic data. For example, we hypothesize that the loss of key hydrogen bond interactions and steric effects lead to conformational changes in interactions of inhibitors with the enzyme binding site. Previous structural studies suggested hydrogen bonds between Val621 and Asp682 of PfPKG and imidazole derivatives ^*12, 25*^ and thiazoles ^*24*^. Our selected docking pose for **3** also captured the importance of engaging Val621 and Asp682 of PfPKG to achieve potency of inhibitors targeting PfPKG’s ATP-binding pocket. We caution that these poses are possibilities predicted via docking studies and not experimentally determined.

We are continuing to develop these models by studying additional compounds in this class and investigate structural changes to the amino pyrrolidine and heterocyclic carboxamide. The initial results reported here furnished design hypotheses for more potent PfPKG inhibitors. As one example, additional sulfone analogs of **3** are being evaluated, along with electronic effects in the isoxazole ring that may influence arene-cation interactions with PfPKG. A second example is the apparent limited steric environment surrounding the tertiary amine in **3**. The conformational rearrangement suggested by this change resulted in a less potent inhibitor and revealed altered interactions within the amine binding site that can be probed with new derivatives. Work is underway to evaluate these hypotheses using crystal structures of one or more of these compounds bound to PfPKG. As new analogs are prepared and evaluated, the docking model can be improved, and its predictive capabilities enhanced.

We envision that if PfPKG inhibitors reach the clinic, they will be used in combination with other anti-malarials to achieve malaria chemoprotection. Combination therapy is essential for reducing the likelihood of parasite resistance. Previous experiments on evolved resistance demonstrated that resistance to ATP-competitive inhibitors of PfPKG is not easily acquired. ^*12, 25*^ Parasites developed only low-level resistance through mutations in proteins other than PfPKGand inhibitor-resistant parasites did not carry the T618Q substitution. ^*25*^ These data suggest that high-level resistance to PfPKG inhibitors via mutations in the target protein is unlikely to be facile and underscore PfPKG’s attractiveness as an anti-malarial drug target.

While our work focuses of ATP-competitive inhibitors of PfPKG, additional chemotypes might be discovered through investigations of allosteric mechanisms of PfPKG activation. The regulatory domain of PfPKG contains three functional cGMP binding sites (an additional one is degenerate and incapable of binding cGMP) compared to two cGMP-binding sites in hPKG. The unique mechanism of PfPKG allosteric activation^*17, 28-30*^ may offer novel means of inhibition - a cGMP analog reduces *in vitro* activity of recombinant PfPKG carrying a single cGMP site.^*30*^ Effect of cGMP analogs on the activity of full-length PfPKG has not yet been examined. Given the urgent need for new chemotypes for an important new target against a disease that causes substantial morbidity and mortality, our approach, of coupling experimental and *in silico* results, and the data presented herein represent a useful advance in PfPKG biochemistry.

## Accession codes

PfPKG: UniProt ID Q8I719

hPKG gene: UniProt ID Q13976

## Acknowledgements

The authors acknowledge support from the National Institutes of Health award R01AI133633-01 to PB, DPR and JJS, the United States Department of Defense award W81XWH2010386 to PB, and the Margaret and Herman Sokol Endowment to JJS and DPR.

## Supporting information

Supplemental information: Supplemental Figures S1-S8 with legends. Spectra for **3, 4**, (proton and carbon NMRs) and **6** (proton NMR).

