## Supplemental Information for "Characterization of novel competitive inhibitors of *P. falciparum* cGMP-dependent protein kinase"

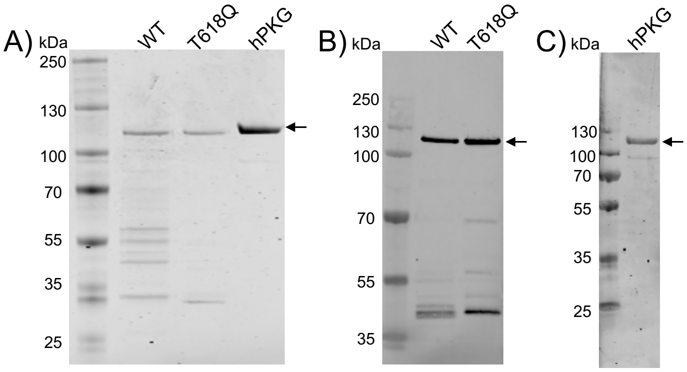


**Figure S1**. **A.** SDS-PAGE of purified and concentrated enzymes used in this study. From left to right: Pre-stained PageRuler Ladder, His-tagged WT, His-tagged T618Q and GST-tagged hPKG. The gel was stained with Instant Blue and visualized on a LiCor CLx. **B.** Western Blot using an anti-His antibody visualized on a LiCor CLx and ImageStudio Lite software. **C.** Western Blot using anti-GST of purified GST-tagged hPKG visualized on a LiCor CLx and ImageStudio Lite software. Arrows indicate full-length proteins.

A)

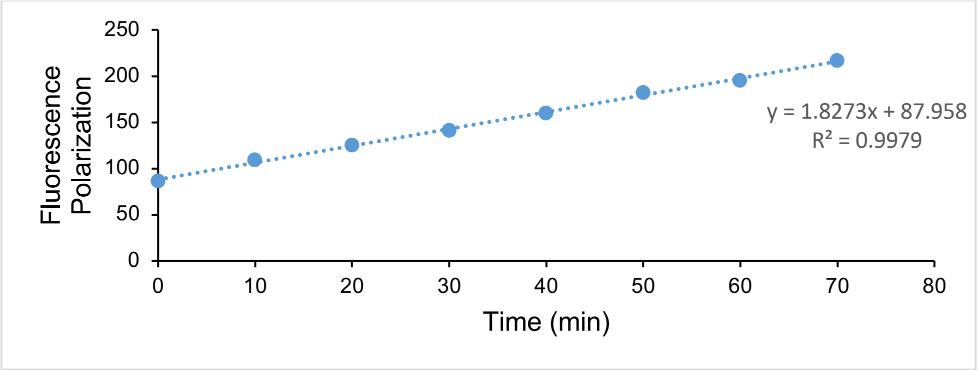


B)

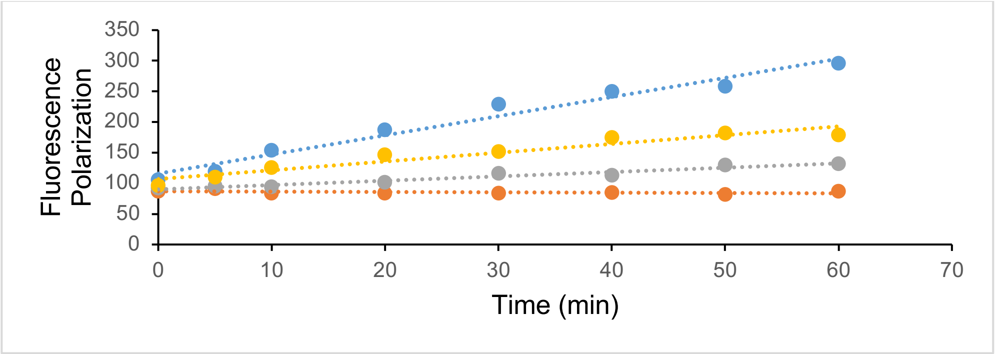


**Figure S2**. **A) Enzymatic activity of WT PfPKG.** A linear relationship between polarization signal and time indicating initial velocity conditions, at 50 µM ATP). **B)** Polarization versus time at other ATP concentrations (orange = 0 µM ATP, grey = 1.5 µM ATP, yellow = 6.25 µM ATP, blue = 100 µM ATP) further demonstrate that rates were collected under initial conditions.

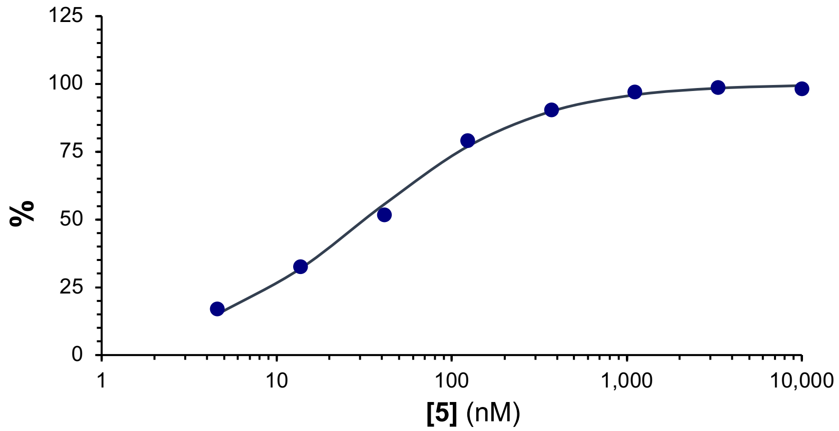


**Figure S3.** Determination of $IC_{50}$ of **5** for WT PfPKG. Data shown are representative of three biological replicates. X-axis is in the log scale. Curve fit was generated using the Solver extension in Microsoft Excel.

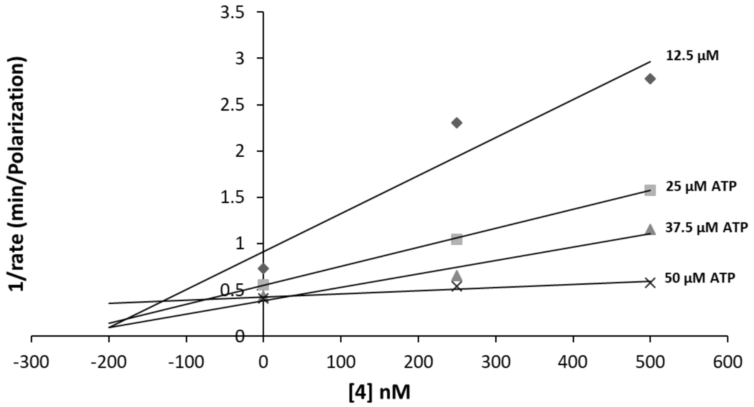


**Figure S4A**: Determination of Ki for **4**. A Dixon Plot representation of observed initial velocities at various concentrations of ATP and **4**. The intersection point corresponds to competitive inhibition and a -Ki of -136 nM. Data shown are representative of three biological replicates.

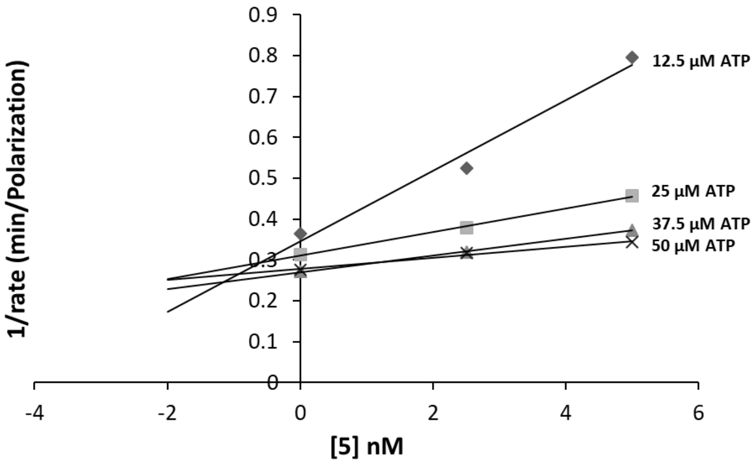


**Figure S4B**: Determination of Ki for **5**. Dixon Plot representation of observed initial velocities at various concentrations of ATP and **5**. The intersection point corresponds to competitive inhibition and a - Ki of -1.4 nM. Data shown are representative of three biological replicates.

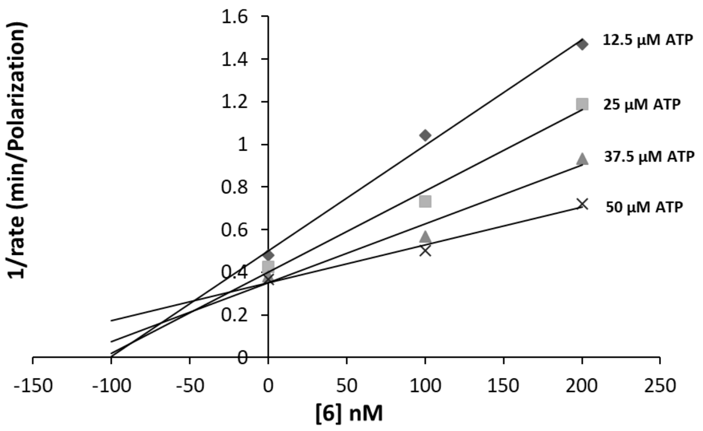


**Figure S4C**: Determination of Ki for **6**. Dixon Plot representation of observed initial velocities at various concentrations of ATP and **6**. The intersection point corresponds to competitive inhibition and a - Ki of -46 nM. Data shown are representative of three biological replicates.

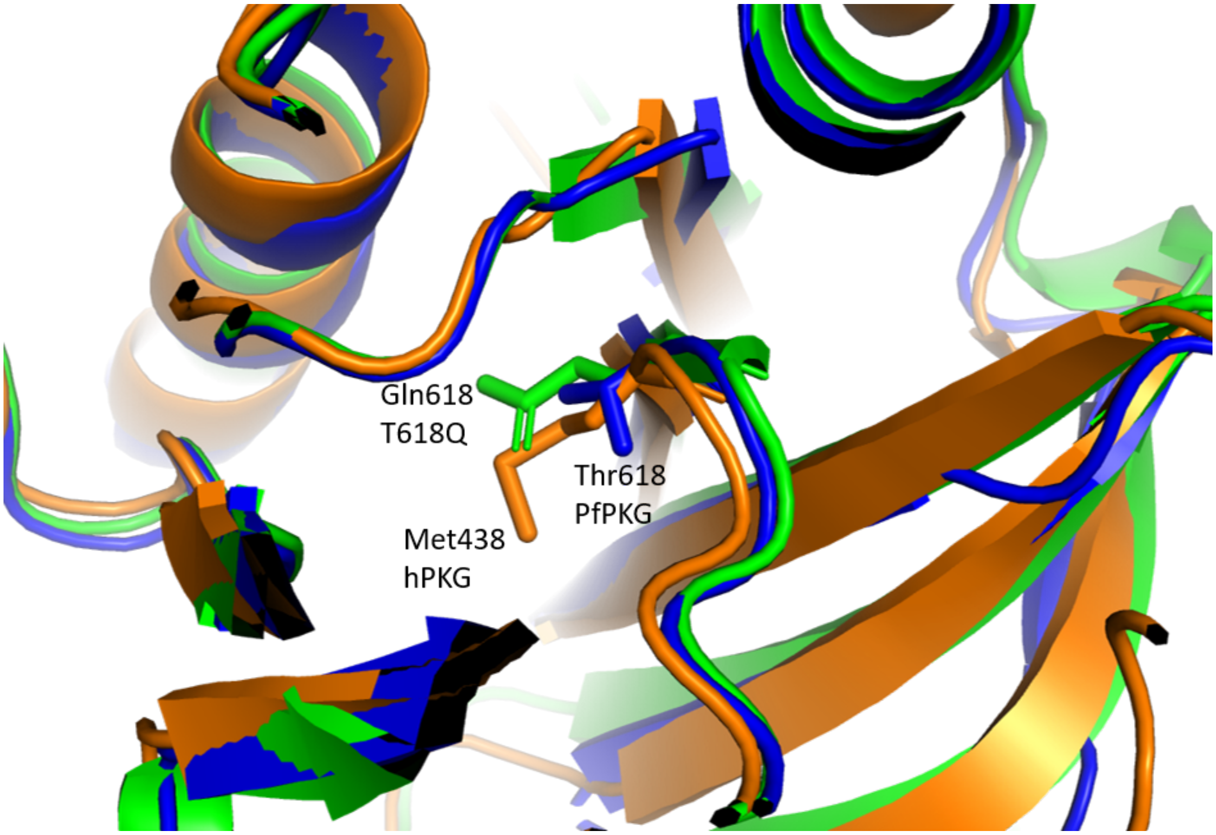


**Figure S5.** 3D alignment PfPKG (blue), T618Q (green) and hPKG (orange). Gatekeeper residues are illustrated as sticks.

```
CLUSTAL format alignment by MAFFT (v7.475)


PfPKG           MEEDDNLKKGNERNKKKAIFSNDDFTGEDSLMEDHLELREKLSEDIDMIKTSLKNNLVCS
MUTANT_PfPKG    MEEDDNLKKGNERNKKKAIFSNDDFTGEDSLMEDHLELREKLSEDIDMIKTSLKNNLVCS
hPKG            ------------------------------------------------------------


PfPKG           TLNDNEILTLSNYMQFFVFKSGNLVIKQGEKGSYFFIINSGKFDVYVNDKKVKTMGKGSS
MUTANT_PfPKG    TLNDNEILTLSNYMQFFVFKSGNLVIKQGEKGSYFFIINSGKFDVYVNDKKVKTMGKGSS
hPKG            -------------------------------------------------------GSG--
                                                                       *.*

PfPKG           FGEAALIHNTQRSATIIAETDGTLWGVQRSTFRATLKQLSNRNFNENRTFIDSVSVFDML
MUTANT_PfPKG    FGEAALIHNTQRSATIIAETDGTLWGVQRSTFRATLKQLSNRNFNENRTFIDSVSVFDML
hPKG            ------------------------------------------------------------


PfPKG           TEAQKNMITNACVIQNFKSGETIVKQGDYGDVLYILKEGKATVYINDEEIRVLEKGSYFG
MUTANT_PfPKG    TEAQKNMITNACVIQNFKSGETIVKQGDYGDVLYILKEGKATVYINDEEIRVLEKGSYFG
hPKG            ------------------------------------------------------------


PfPKG           ERALLYDEPRSATIIAKEPTACASICRKLLNIVLGNLQVVLFRNIMTEALQQSEIFKQFS
MUTANT_PfPKG    ERALLYDEPRSATIIAKEPTACASICRKLLNIVLGNLQVVLFRNIMTEALQQSEIFKQFS
hPKG            ------------------------------------------------------------


PfPKG           GDQLNDLADTAIVRDYPANYNILHKDKVKSVKYIIVLEGKVELFLDDTSIGILSRGMSFG
MUTANT_PfPKG    GDQLNDLADTAIVRDYPANYNILHKDKVKSVKYIIVLEGKVELFLDDTSIGILSRGMSFG
hPKG            --------------------------------------------LDDVS-----------
                                                            ***.*

PfPKG           DQYVLNQKQPFKHTIKSLEVCKIALITETCLADCLGNNNIDASIDYNNKKSIIKKMYIFR
MUTANT_PfPKG    DQYVLNQKQPFKHTIKSLEVCKIALITETCLADCLGNNNIDASIDYNNKKSIIKKMYIFR
hPKG            -----------------------------------------------NKA----------
                                                               **

PfPKG           YLTDKQCNLLIEAFRTTRYEEGDYIIQEGEVGSRFYIIKNGEVEIVKNKKRLRTLGKNDY
MUTANT_PfPKG    YLTDKQCNLLIEAFRTTRYEEGDYIIQEGEVGSRFYIIKNGEVEIVKNKKRLRTLGKNDY
hPKG            ------------------YEDAE--------------------------------AKAKY
                                  **:.:                                .* .*

PfPKG           FGERALLYDEPRTASVISKVNNVECWFVDKSVFLQIIQGPMLAHLEERIKMQDTKVEMDE
MUTANT_PfPKG    FGERALLYDEPRTASVISKVNNVECWFVDKSVFLQIIQGPMLAHLEERIKMQDTKVEMDE
hPKG            EAEAAFF-----------------------------------------------ANLKLSD
                 .* *::                                              :::::.:

PfPKG           LETERIIGRGTFGTVKLVHHKPTKIR-YALKCVSKRSIINLNQQNNIKLEREITAENDHP
MUTANT_PfPKG    LETERIIGRGTFGTVKLVHHKPTKIR-YALKCVSKRSIINLNQQNNIKLEREITAENDHP
hPKG            FNIIDTLGVGGFGRVELVQLKSEESKTFAMKILKKRHIVDTRQQEHIRSEKQIMQGAHSD
                ::    :* * ** *:**: *. : : :*:* :.** *:: .**::*: *::*     .

PfPKG           FIIRLVRTFKDSKYFYFLTELVTGGELYDAIRKLGLLSKSQAQFYLGSIILAIEYLHERN
MUTANT_PfPKG    FIIRLVRTFKDSKYFYFLQELVTGGELYDAIRKLGLLSKSQAQFYLGSIILAIEYLHERN
hPKG            FIVRLYRTFKDSKYLYMLMEACLGGELWTILRDRGSFEDSTTRFYTACVVEAFAYLHSKG
                **:** ********:*:**    ****:  :*. * :..* ::** ..:: *: ***.:.

PfPKG           IVYRDLKPENILLDKQGYVKLIDFGCAKKV--QGRAYTLVGTPHYMAPEVILGKGYGCTV
MUTANT_PfPKG    IVYRDLKPENILLDKQGYVKLIDFGCAKKV--QGRAYTLVGTPHYMAPEVILGKGYGCTV
hPKG            IIYRDLKPENLILDHRGYAKLVDFGFAKKIGFGKKTWTFCGTPEYVAPEIILNKGHDISA
                *:********::**::**.**:*** ***:     :::*: ***.*:***:**.**:. :.

PfPKG           DIWALGICLYEFICGPLPFGNDEEDQLEIFRDILTG--QLTFPDYVTDTDSINLMKRLLC
MUTANT_PfPKG    DIWALGICLYEFICGPLPFGNDEEDQLEIFRDILTG--QLTFPDYVTDTDSINLMKRLLC
hPKG            DYWSLGILMYELLTGSPPFSG--PDPMKTYNIILRGIDMIEFPKKIA-KNAANLIKKLCR
                * *:*** :**:: *. **..    * :: :. ** *    : **. :: .:: **:*:*

PfPKG           RLPQGRIGCSINGFKDIKDHPFFSNFNWDKLAGRLLDPPLVSKSETYAEDIDIKQIEEED
MUTANT_PfPKG    RLPQGRIGCSINGFKDIKDHPFFSNFNWDKLAGRLLDPPLVSKSETYAEDIDIKQIEEED
hPKG            DNPSERLGNLKNGVKDIQKHKWFEGFNWEGLRKGTLTPPIIP---SVASPTDTSNFDSFP
                  *. *:*   **.***:.* :*..***: *    * **::.   : *.  * .:::.

PfPKG           AEDDEEPLNDEDNWDIDF
MUTANT_PfPKG    AEDDEEPLNDEDNWDIDF
hPKG            EDNDEPPPDDNSGWDIDF
                 ::** * :*:..*****
```

**Figure S6.** Sequence alignment PfPKG, T618Q and hPKG. Gatekeeper residues are highlighted in red dashes.

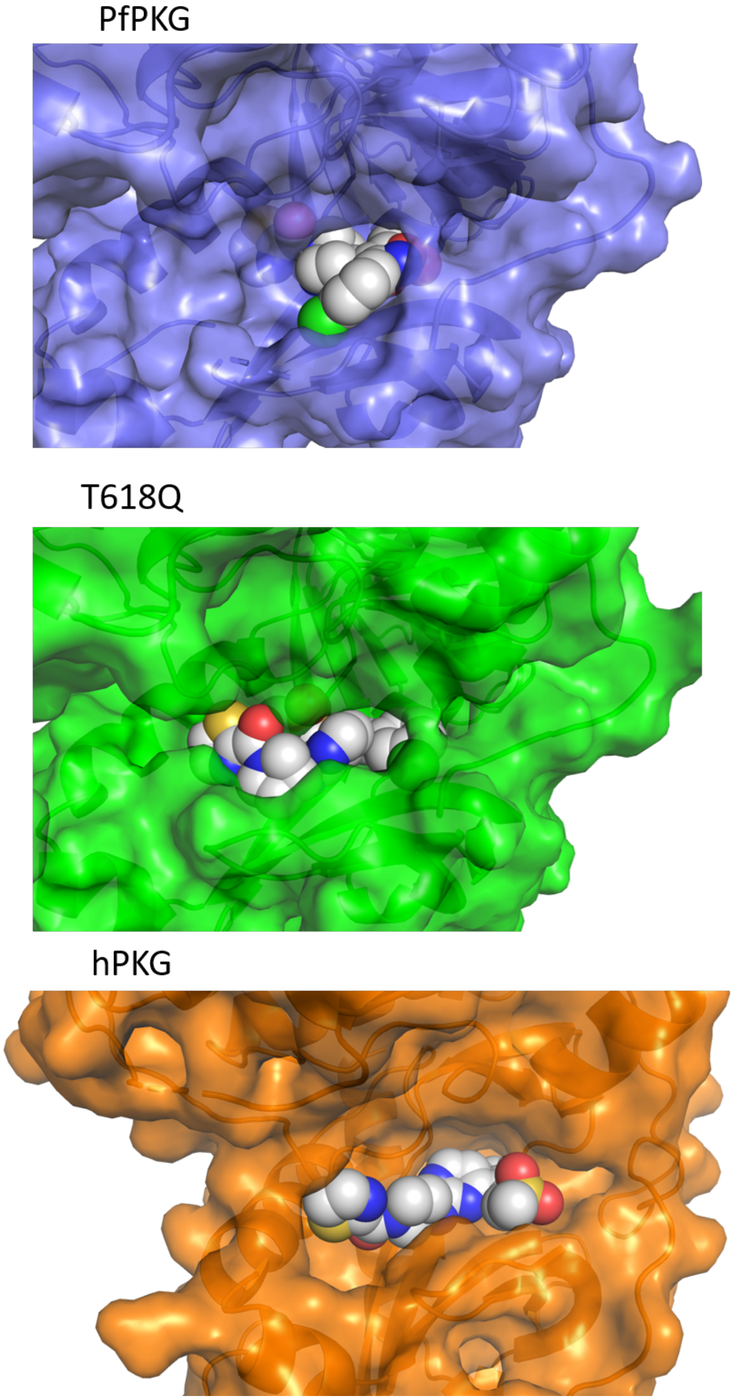


**Figure S7.** Cartoons illustrating differences in binding pocket size: Surface representations of and PfPKG (blue), T618Q (green) and hPKG (orange) and **3** as spheres.

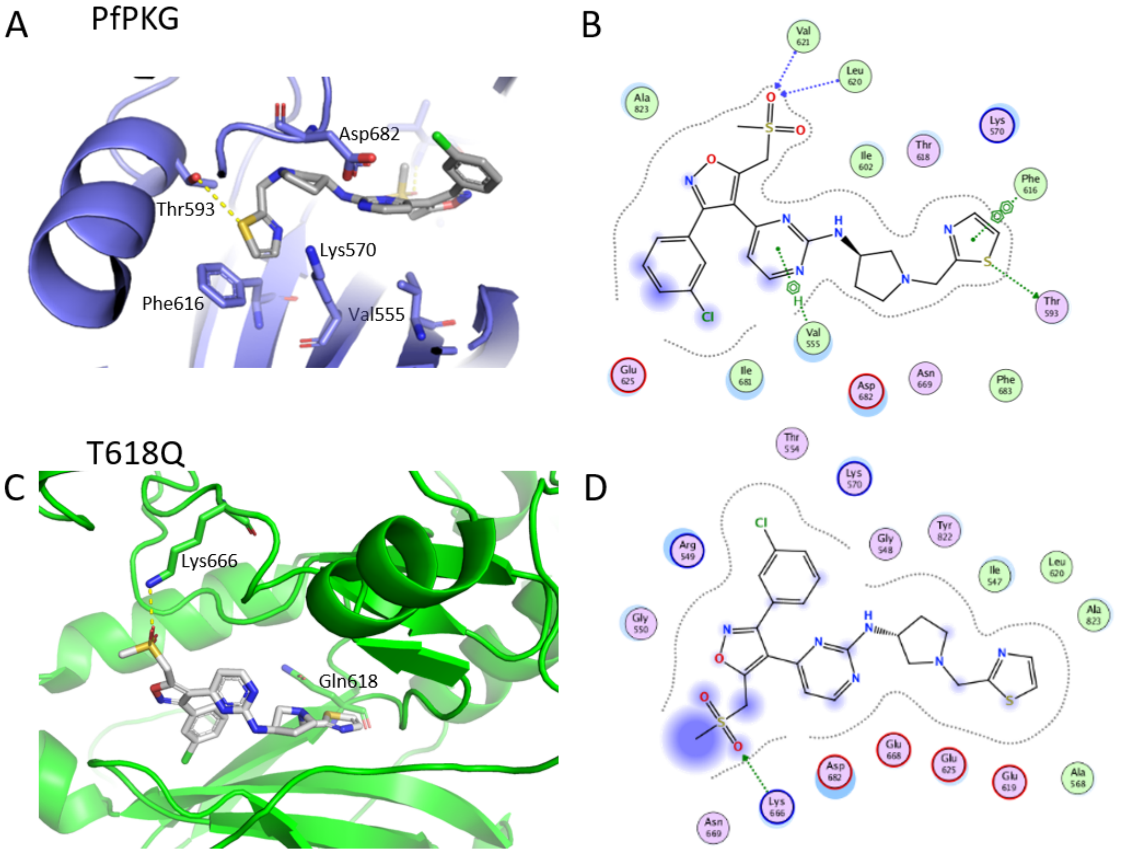


**Figure S8.** Molecular docking of **4** into PfPKG (A, B) and PfPKG T618Q mutant (C, D). Substitution of T618, located centrally in the binding pocket, to Q618 changes reduces the size of the binding pocket and changes the landscape of interactions with **4**. A) In WT PfPKG, **4** interacts with Val621, Leu620, Phe616, Thr693, Val555. B) In T618Q, **4** interacts only with Lys666. Docking of **4** predicts greater solvent exposure in T618Q than in WT. These predictions are consistent with the activity of **4** against WT and T618Q enzymes. Figures A, C and E were prepared using PyMOL. Figures B, D and F were prepared using MOE.

"MSU-RY-1-102 (CH2SO2Me-2-Thiazole)"  1  1  C:\Bruker\TopSpin3.2\data\Rotella\Rammohan

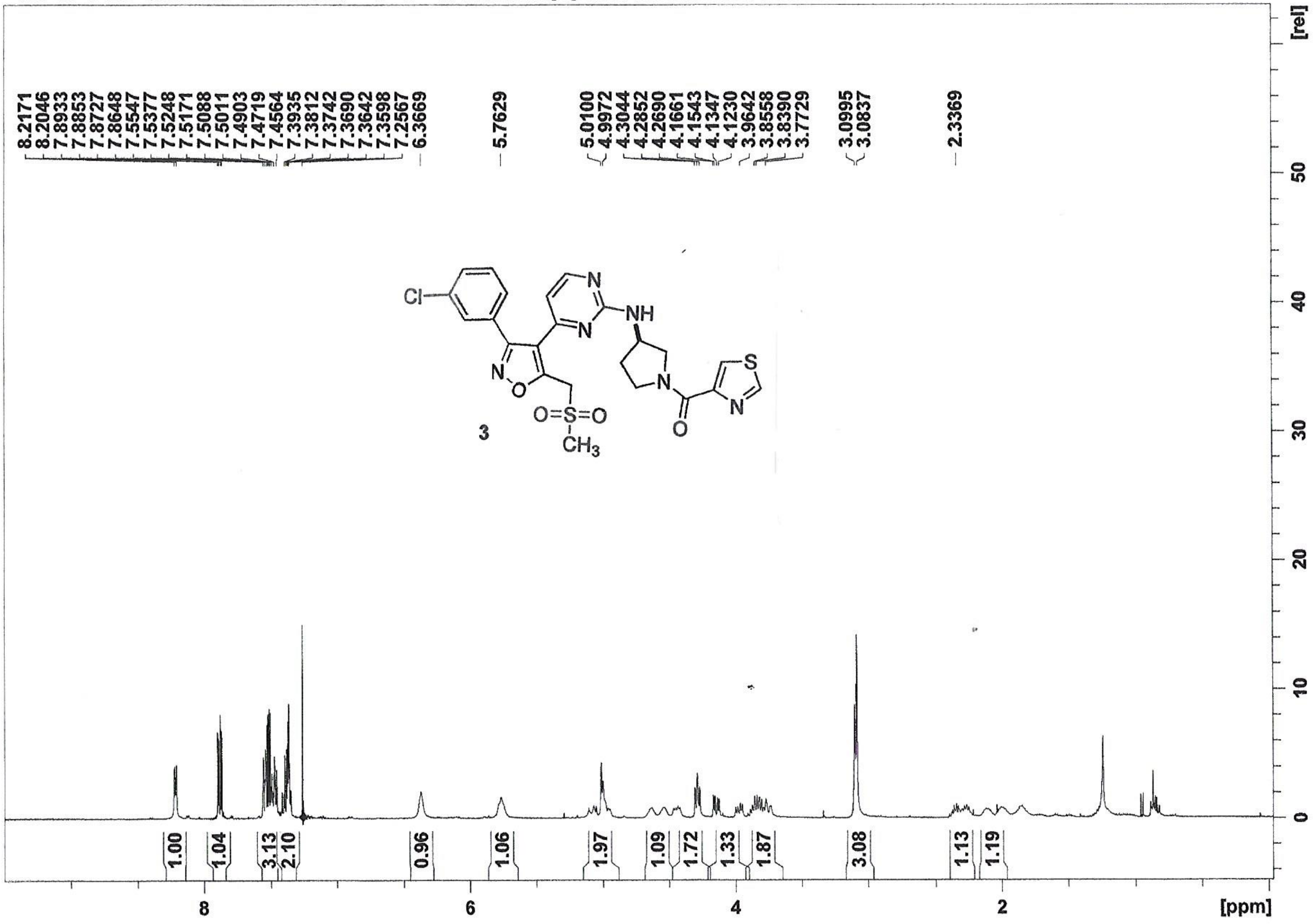

"MSU-RY-1-102 13C"  13  1  C:\Bruker\TopSpin3.2\data\Rotella\Rammohan

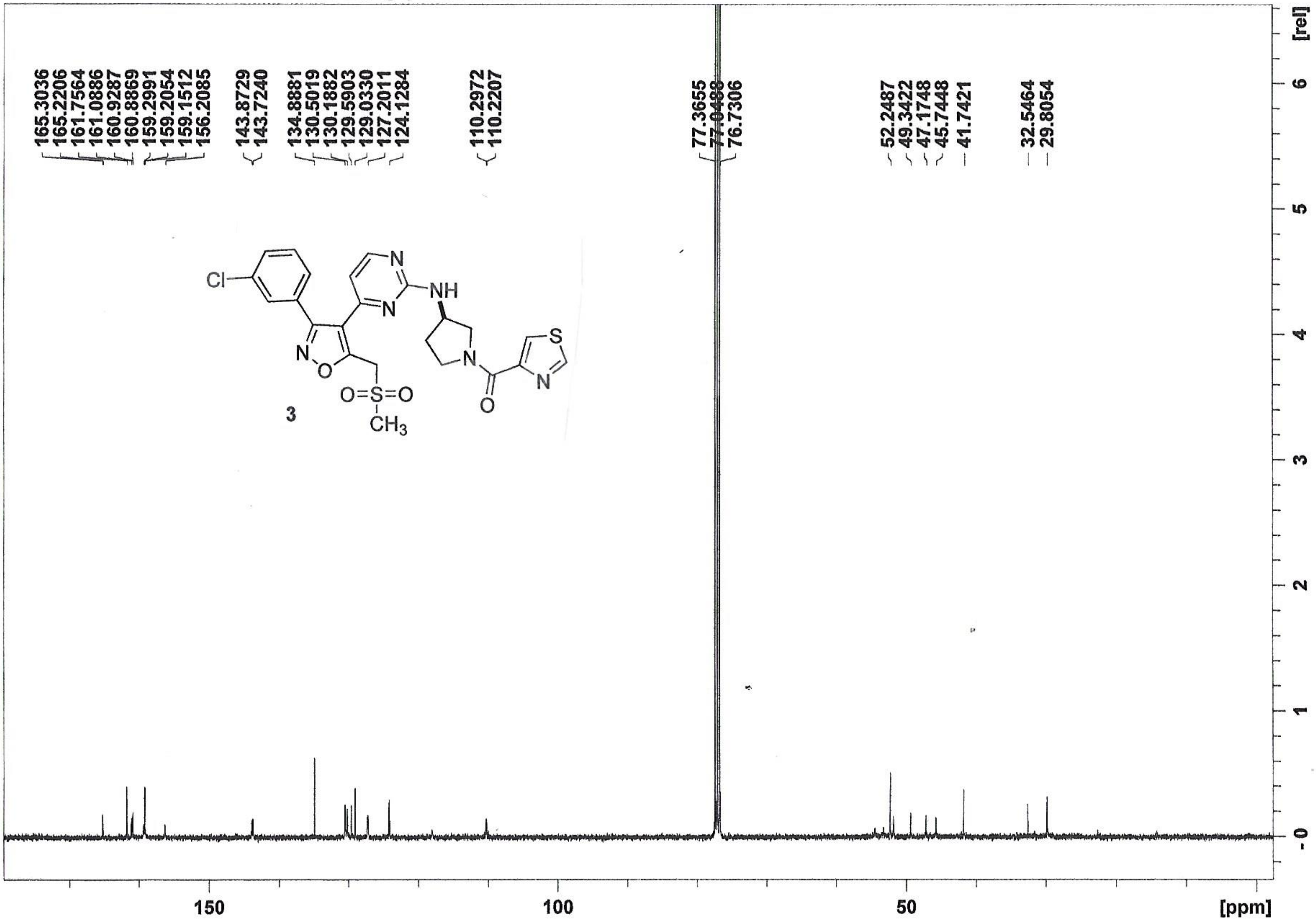

"MSU-RY-1-113 (CH2SO2Me-Me-2-Thiazole)"  1  1  C:\Bruker\TopSpin3.2\data\Rotella\Rammohan

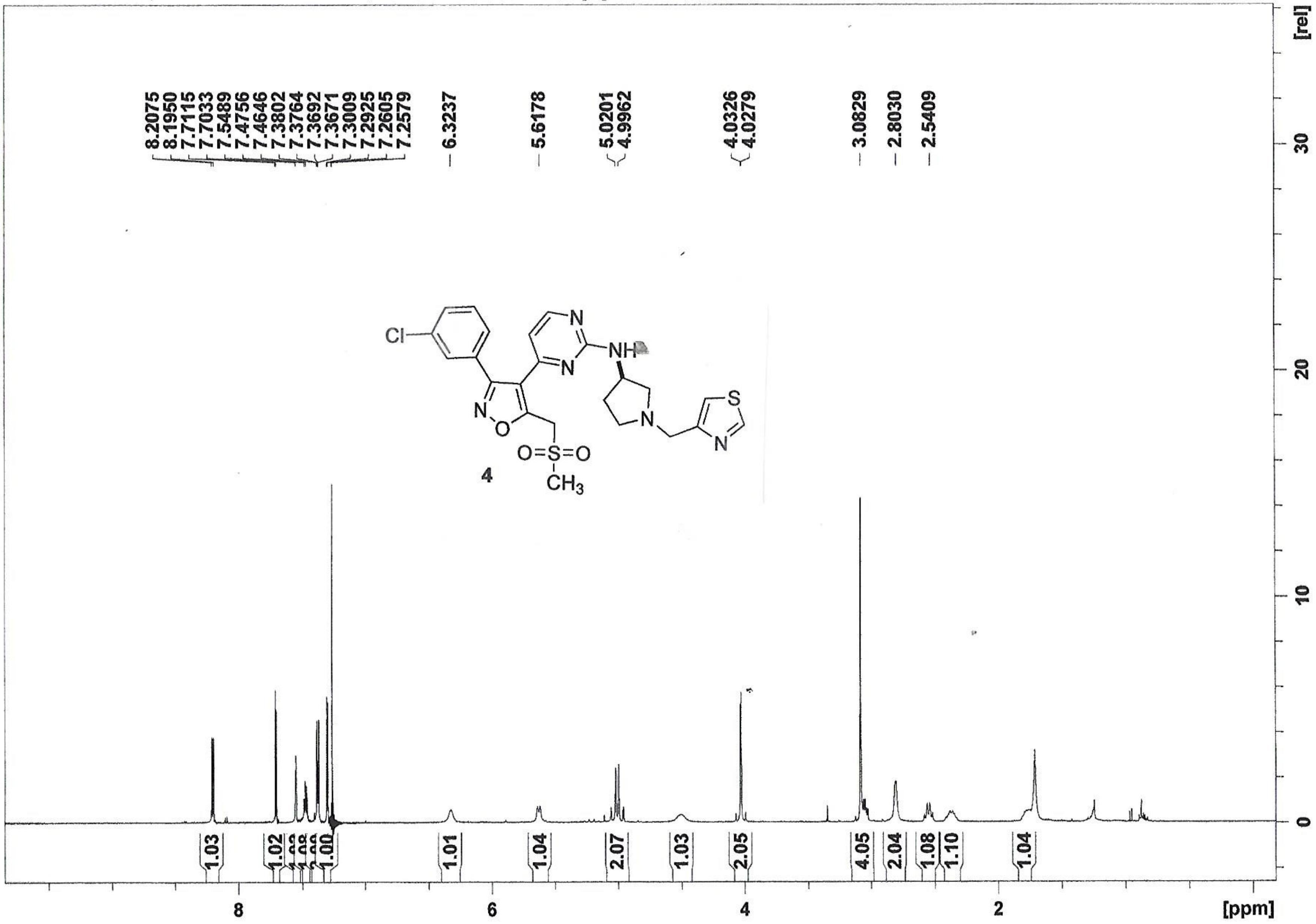

"MSU-RY-1-113 13C"  13  1  C:\Bruker\TopSpin3.2\data\Rotella\Rammohan

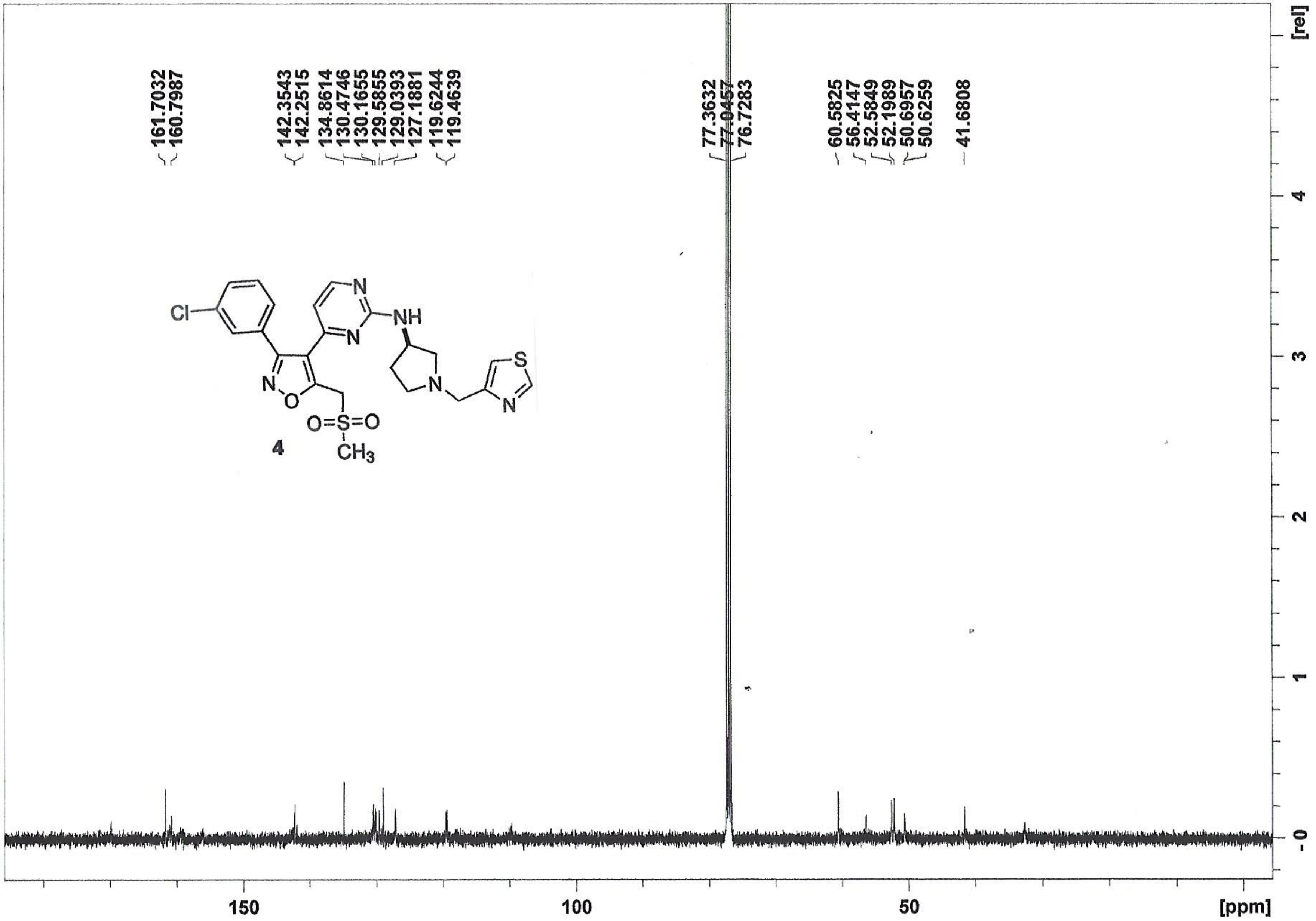

MSU-RTC-1-071-1 1 1 C:\Bruker\TopSpin3.2\data\Rotella\Rammohan

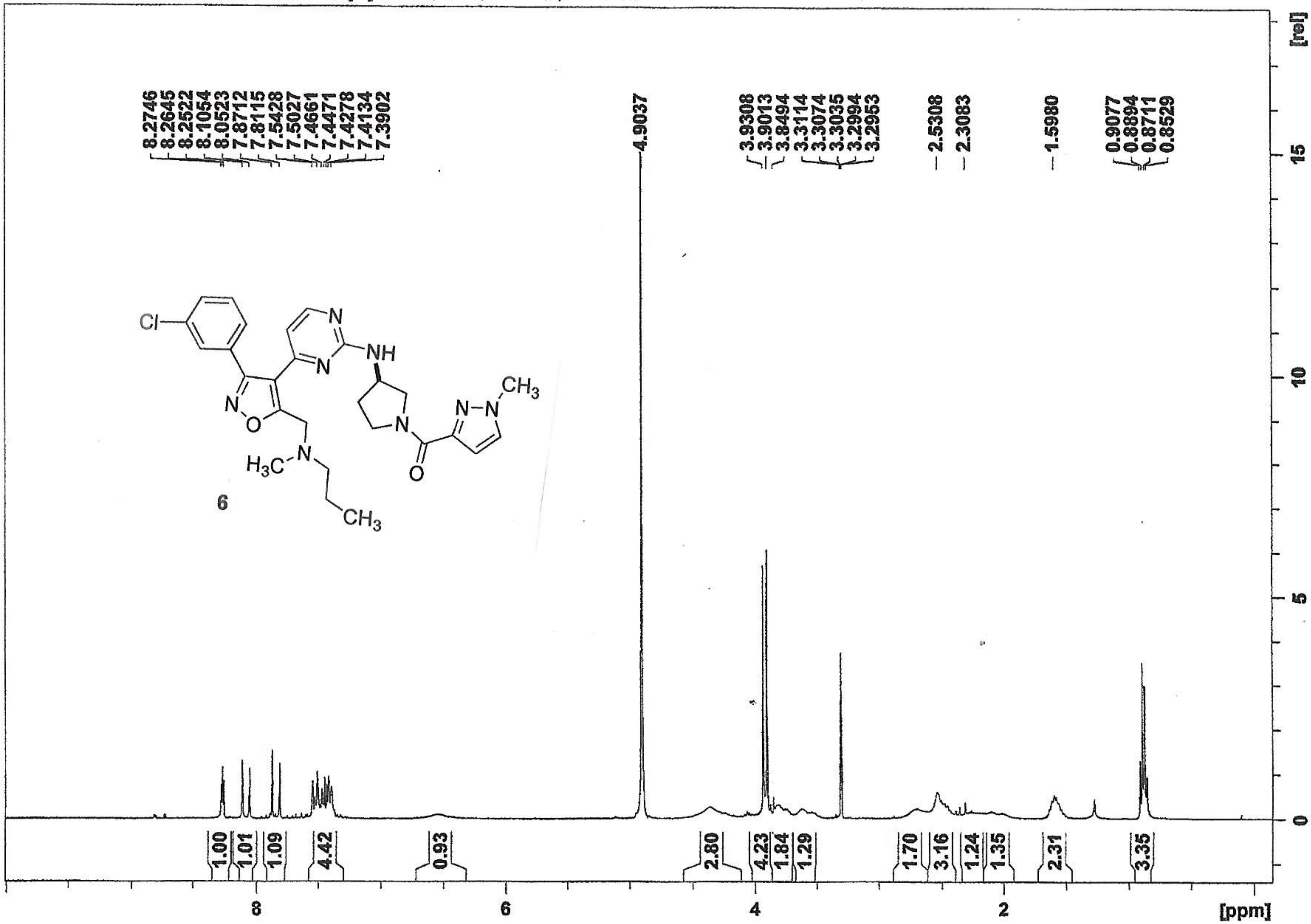


6
